# Microbial interactions affect the tempo and mode of antibiotic resistance evolution

**DOI:** 10.1101/2024.06.06.597700

**Authors:** Laurens E. Zandbergen, Joost van den Heuvel, Andrew D. Farr, Bas J. Zwaan, J. Arjan G. M. de Visser, Marjon G. J. de Vos

## Abstract

The global rise of antibiotic resistance impedes the treatment of bacterial infections. To limit the emergence and evolution of antibiotic resistance it is important to understand how bacterial interactions in multispecies communities affect the course of evolution. We investigated how ecological interactions between microbes derived from polymicrobial urinary tract infections affect the tempo and mode of antibiotic resistance evolution. We show that for representative strains of three uropathogens, *Escherichia coli, Klebsiella pneumoniae* and *Enterococcus faecium,* the rate and evolutionary trajectories towards antibiotic resistance differ depending on the conditioned medium mediated interactions with other microbes that alter their growth and antibiotic tolerance. Replicate lineages of the same species evolved under similar ecological conditions show parallel evolutionary trajectories, and resistance mutations and other functional targets selected differed between these conditions. Our findings demonstrate that bacterial interactions differentially affect the evolutionary potential of antibiotic resistance evolution.

## Introduction

Bacterial pathogens often live in multispecies communities. In such microbial communities, bacteria interact, which affects their growth, survival and coexistence (1–5). Bacterial infections are often treated with antibiotics, whose efficacy may also depend on ecological interactions with community members (6). For instance, co-existing bacterial species in pathogenic communities can mediate the growth and survival of community members in the presence of antibiotics (7–9) through antibiotic degradation (10,11), change the metabolic landscape affecting pathogen fitness (12–14), lower antimicrobial efficacy or alter the transport of antibiotics (7,15), or change the induction of signal molecules that induce resistance mechanisms (16).

Ecological interactions can therefore affect the evolvability of antibiotic resistance in several ways. Firstly, they may alter the mutation supply, by affecting the population size or mutation rate (17–20). Additionally, ecological interactions may alter the fitness effect of mutations, mediating the selective accessibility of adaptive mutations, as genetic and environmental factors are known to modulate the effect of mutations (21–23) and therefore potentially morph the shape of the adaptive landscape (24,25).

In clinical practice, urinary tract infections (UTIs) are treated with antibiotics. Recurrence of infection after the discontinuation of the antimicrobial therapy in treated patients can be high (up to 50% in the elderly population) (26). This is not surprising, as infections are often polymicrobial and many UTI isolates are resistant to commonly prescribed antibiotics (1,27,28). To investigate whether different microbial interactions have an effect on the evolution of antibiotic resistance, we evolved three focal species (*Escherichia coli, Klebsiella pneumoniae* and *Enterococcus faecium*) in the conditioned medium of several species (*K. pneumoniae*, *E. coli*, *E. faecium*, *Staphylococcus haemolyticus* and *Pseudomonas aeruginosa*). The bacteria were isolated from elderly patients diagnosed with polymicrobial urinary tract infections (1). These pathogens are part of the so-called ESKAPEE bacteria, which are known for their ability to evade antibiotics due to their increasing multi-drug resistance (29). We hypothesize that ecological interactions between uropathogens determine the evolutionary potential of antibiotic resistance evolution through the modulation of population sizes and changing the shape of the adaptive landscape.

## Materials and Methods

### Bacterial isolates and growth medium

The five uropathogen isolates used in this study were previously isolated anonymously from different elderly patients (>70 years old) suffering from urinary tract infection (1). These five isolates (*E. coli*, *K. pneumonia*, *E. faecium*, *S. haemolyticus*, and *P. aeruginosa*) were selected based on their consistent planktonic growth in Artificial Urine Medium (AUM) and for their observed growth affecting interactions in a previous study (2,6). Isolates were cultured in Artificial Urine Medium (AUM) (30).

### Preparation of conditioned medium

Spent medium of *E. coli*, *K. pneumonia*, *E. faecium*, *S. haemolyticus*, and *P. aeruginosa* isolates was prepared by culturing in AUM, followed by centrifugation and filtering. To prepare conditioned medium, the AUM spent medium fraction was replenished with AUM constituents.

### Serial dilution experimental evolution protocol

*K. pneumoniae* and *E. faecium* were evolved for 30 bi-daily transfers, *E. coli* was evolved for 35 bi-daily transfers. Focal species were cultured in 5 ml preheated conditioned medium, in 15 ml screw cap falcon tubes, with the caps slightly opened. Each transfer *K. pneumoniae* and *E. coli* were diluted 500 times, and *E. faecium* was diluted 100 times, due to its inherent smaller population size in the artificial urine medium and urine. *K. pneumoniae* and *E. coli* were evolved in the conditioned media of *K. pneumoniae*, *E. coli*, *E. faecium*, *Staphylococcus haemolyticus. E. faecium* was additionally also evolved in *P. aeruginosa* conditioned medium. Each focal isolate-conditioned medium combination was replicated five times. The Gram-negative isolates *K. pneumoniae* and *E. coli* were evolved in trimethoprim-sulfamethoxazole (ratio 1:5, as in clinical use (31)) and *E. faecium* in vancomycin (32). The level of antibiotics at the start of the experiments was set at a 50% sub-MIC concentration, 0.05 and 0.0125 µg/mL trimethoprim for *K. pneumoniae,* and *E. coli* respectively, and 0.25 µg/mL vancomycin for *E. faecium*. If the cultures exceeded a population size threshold (for *K. pneumoniae* and *E. coli* 0.1 OD600 and 0.05 OD600 for *E. faecium)*, inferred through an optical density assessment, similar to mcFarland standard assay (33), the antibiotic concentration was increased by 20%, otherwise the antibiotic concentration was kept at the same level as the previous transfer.

### Growth Measurements

Three clones of three of the five evolved replicate lineages of each focal species, evolved in each conditioned medium, and the wild types of each focal species were tested in phenotypic growth assays. Growth rates were obtained by incubation in a BMG Clariostar or Fluorostar microplate reader at 37°C under aerobic conditions and shaken continuously at 500 rpm. The growth rate was defined by the shortest obtained doubling time and quantified by the steepest linear fit of the exponential phase of log(OD600). Population density was inferred from OD600 absorbance measurements as well, taking the average for all the measurements of the stationary phase of the growth curve.

### Antibiotic tolerance assays

Growth measurements were taken of wild type isolates of *K. pneumoniae*, *E. faecium* and *E. coli* were inoculated with 2 µl of overnight cultures in triplicates in conditioned medium and AUM reference medium in a concentration gradient with two-fold increases in antibiotic concentration in 96-well plates. Plates were incubated for 24h at 37°C. *K. pneumoniae* and *E. coli* were assayed in the presence of trimethoprim-sulfamethoxazole and *E. faecium* in the presence of vancomycin. The fold change tolerance was calculated by dividing the concentration that limited growth in a particular conditioned medium by the concentration of antibiotic that limited growth in AUM.

### MIC assays

Starting Minimum Inhibitory Concentration (MIC) of the wild-types were measured in a serial dilution assay of the desired antibiotics in AUM. 200 µL cultures were created in a 96-well plate including a two-fold serial dilution of the desired antibiotic (vancomycin for *E. faecium* and a 1:5 mixture of trimethoprim and sulfamethoxazole for both *E. coli* and *K. pneumoniae*) ranging from 0.125 to 1 µg/mL for the vancomycin and 0.00625 to 0.2 µg/mL for trimethoprim:sulfamethoxazole, in triplicate. Cultures without antibiotics were used as control. Cultures were incubated over 24h at 37°C at 200 rpm shaking conditions. Whether or not bacteria had visibly grown was determined by measuring OD600 and subtracting AUM without bacteria as blank. OD600 values <0.05 were inspected visually to determine complete inhibition. MIC of final transfer mutants were determined on agar using a modified version of the TDtest described previously by Gefen et al. (34), Supplementary Text 1.

### DNA extraction

Cultures of evolved clones were grown overnight in 5ml LB, cells were pelleted, and DNA was extracted using a Maxwell 16 Instrument in combination with the Maxwell® 16 Tissue LEV Total RNA Purification Kit Custom (XAS 1220) (Promega, Madison, WI, USA) following manufacturer’s instructions. The DNA was eluted in 30 µL DNAse free water, and quantified using Nanodrop 2000c Spectrophotometer (Thermo Fisher Scientific, USA) and gel electrophoresis.

### Sequencing

Three clones of three of the five evolved replicate lineages of each focal species, evolved in each conditioned medium, were prepared for sequencing. DNA was extracted using the Maxwell® 16 Tissue LE Total RNA purification kit (article number XAS1220, specifically tailored towards DNA extraction). Sequencing was performed at Baseclear BV Leiden. Of the ancestor and the evolved clones, paired-end 125 bp reads were generated using the Illumina HiSeq2500 or MiSeq system. Of the ancestors, long sequence reads were generated using the Oxford Nanopore GridION X5 system to facility the mapping of Illumina reads of the evolved mutants, and ancestor.

### dN/dS

We calculated the dN/dS ratio of all mutations in the genomes of each sequenced end point clone, by dividing the number of non-synonymous mutations by the number of synonymous mutations, and controlling for mutation bias (35).

### Calculation *H*-index

To quantify the repeatability of genomic evolution, the *H*-index previously described by Schenk et al. (17) was used to estimate the pairwise fraction of shared mutations. The *H-*index was calculated at the function (COG) and gene level. All mutants were compared pairwise, calculated using (17):

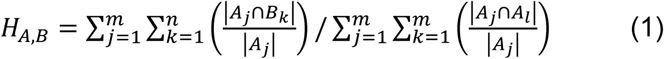

Which compares genotype *A* and *B* with *m* and *n* mutations (with index *j* and *k*), respectively. Here, *A_j_* ∩ *A_l_* denotes the mutations that genotypes *A* and *B* have in common. The *H*-index is calculated for both genotypes separately (*H_A,B_* and *H_B,A_*) to take into account asymmetries in overlap between mutations of different sequence size and compared using (17):

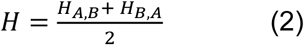

The final *H-*index value would fall between 0 (no repeated evolution) and 1 (complete parallelism). When the *H*-index is 0.5, it indicates that the average of the two compared mutants have half of their mutations shared between them.

### Mutation rate assay

To estimate the intrinsic mutation rate of *E. faecium* and *K. pneumoniae*, we used a high-throughput test which is similar to the fluctuation test developed by Luria and Delbrück (36).

### Statistical analysis

For each genotype, measurements (of phenotypes such as on growth rate, population size, resistance or tolerance levels) were performed in either triplicates or quadruplicates. The null hypotheses were rejected if the *P*-value obtained from the test was lower than a significance level of 0.05. For statistical tests with multiple pairwise comparisons, *P*-values were adjusted using the Holm-Bonferroni method (37).

Additional information on all Materials and Methods can be found in Supplementary text 1.

## Results

### Conditioned media promote the adaptive radiation of antibiotic resistance evolution

To investigate the differential effect of microbial interactions on the evolution of antibiotic resistance, we evolved single isolates of focal species *K. pneumoniae*, *E. coli* and *E. faecium* by bi-daily serial transfers in conditioned artificial urine medium (AUM) supplemented with antibiotics. Conditioned medium was used as a proxy for microbial interactions. It was prepared by creating spent media of cultures grown in AUM followed by replenishment of the medium with fresh nutrients (Materials and Methods) (6). When the evolving isolates had reached a predetermined population size threshold at the moment of transfer, the antibiotic concentration was increased by 20% (Materials and Methods). Each combination of focal isolate and conditioned medium was replicated five times. The two Gram-negative species, *K. pneumoniae* and *E. coli*, were evolved in the presence of trimethoprim-sulfamethoxazole (ratio 1:5, similar to clinical use (31)), which inhibits DNA synthesis (38). *E. faecium* was evolved in the presence of the Gram-positive specific antibiotic vancomycin, which inhibits cell wall synthesis (39); initial antibiotic concentrations were 50% of the minimal inhibitory concentration (MIC) in all cases. *E. coli* was evolved over 70 days (35 transfers) and *K. pneumoniae* and *E. faecium* for 60 days (30 transfers) (Materials and Methods).

We find that conditioned media differentially affect the rate of evolution and final level of antibiotics in the environment (Figure 1). Notably, the evolutionary trajectories of the replicates per conditioned medium are highly parallel. This is especially the case for *K. pneumoniae* and *E. faecium* (Figure 1B,C). Comparing the rate of evolution defined as the slope of the graphs and the final antibiotic concentration reached per focal species and conditioned medium, we find significant differences for all conditioned medium for *E. faecium* (*P<*0.05 for all pairwise comparisons, Welch’s t-test with Holm-Bonferroni correction). For *K. pneumoniae* we find significant differences (*P<*0.05, evolutionary rate and final antibiotic concentration) for all conditioned media, except for the overlapping trajectories of *E. coli* and *S. haemolyticus* conditioned media. The evolutionary trajectories of *E. coli* per conditioned medium environment also display parallelism (Figure 1A), but are more variable compared to *K. pneumoniae* and *E. faecium* (Figure 1B,C); we find significant differences for all conditioned media between conditions (*P*<0.05, evolutionary rate and final antibiotic concentration), except for the partially overlapping trajectories of *S. haemolyticus* and *E. faecium,* and *S. haemolyticus and E. coli* conditioned medium (*P*=0.059 and *P*=0.10, respectively). This shows that the ecological interactions affect the course of antibiotic resistance evolution.

**Figure 1.**
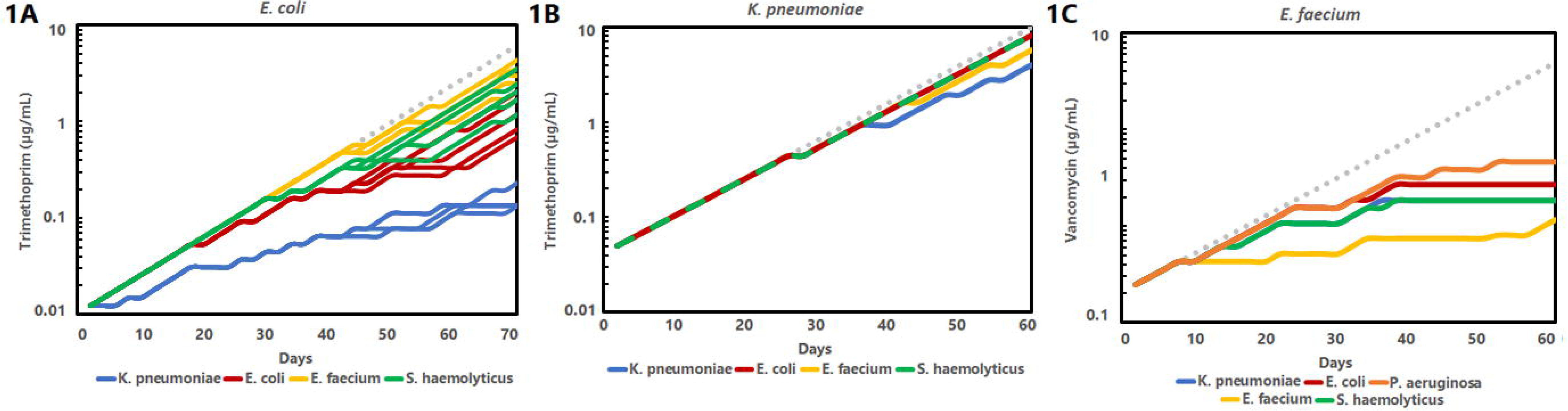
Evolutionary trajectories of uropathogenic bacteria in conditioned media. The gray dotted graph in each figure represents the maximum antibiotic concentration possible in the experimental setup per transfer. **A,** The phenotypic trajectories of *E. coli* evolution in the presence of sulfamethoxazole-trimethoprim, with ratio sulfamethoxazole:trimethoprim 5:1. *E. coli* was evolved in the presence of the conditioned media of *K. pneumoniae, E. coli, E. faecium* and *S. haemolyticus*. The five replicates evolved along similar trajectories, but are more divergent than those of *K. pneumoniae* and *E. faecium*. **B,** The phenotypic trajectories of *K. pneumoniae* evolution in the presence of sulfamethoxazole-trimethoprim. Ratio sulfamethoxazole:trimethoprim is 5:1. *K. pneumoniae* was evolved in the presence of the conditioned media of *K. pneumoniae, E. coli, E. faecium* and *S. haemolyticus*. Five replicates were evolved in each conditioned medium and all replicate trajectories overlapped. **C,** The phenotypic trajectories of *E. faecium* evolution in the presence of vancomycin. *E. faecium* was evolved in the presence of the same conditioned media as *K. pneumoniae* and *E. coli*, and also in the conditioned medium of *P. aeruginosa*. The trajectories of all five replicates per conditioned medium overlapped.

### Response to selection correlates with ecological effects on population size

Bacterial interactions, mediated by direct interactions or conditioned medium, are known to affect the population sizes, for instance by competition for nutrients or cross-feeding. To investigate whether population size effects due to bacterial interactions are related to the level of evolution observed, we correlate the maximum population sizes, as inferred by OD600 measurements, of the ancestors in conditioned medium, with the final antibiotic levels tolerated in the evolution experiments (Figure 2A-C). Indeed, the antibiotic concentration at the final transfer correlates significantly with the population size in conditioned medium for all three focal species. We find a strong positive correlation for *E. coli* and *E. faecium*, and a moderate correlation for *K. pneumoniae* (*R^2^*=0.86, 0.93 and 0.45, for *E. coli*, *E. faecium* and *K. pneumoniae* resp., *P*<0.001 for *E. coli* (*N=*20) and *E. faecium* (*N*=25), and *P*=0.017 for *K. pneumoniae* (*N*=20), Figure 2A-C).

**Figure 2.**
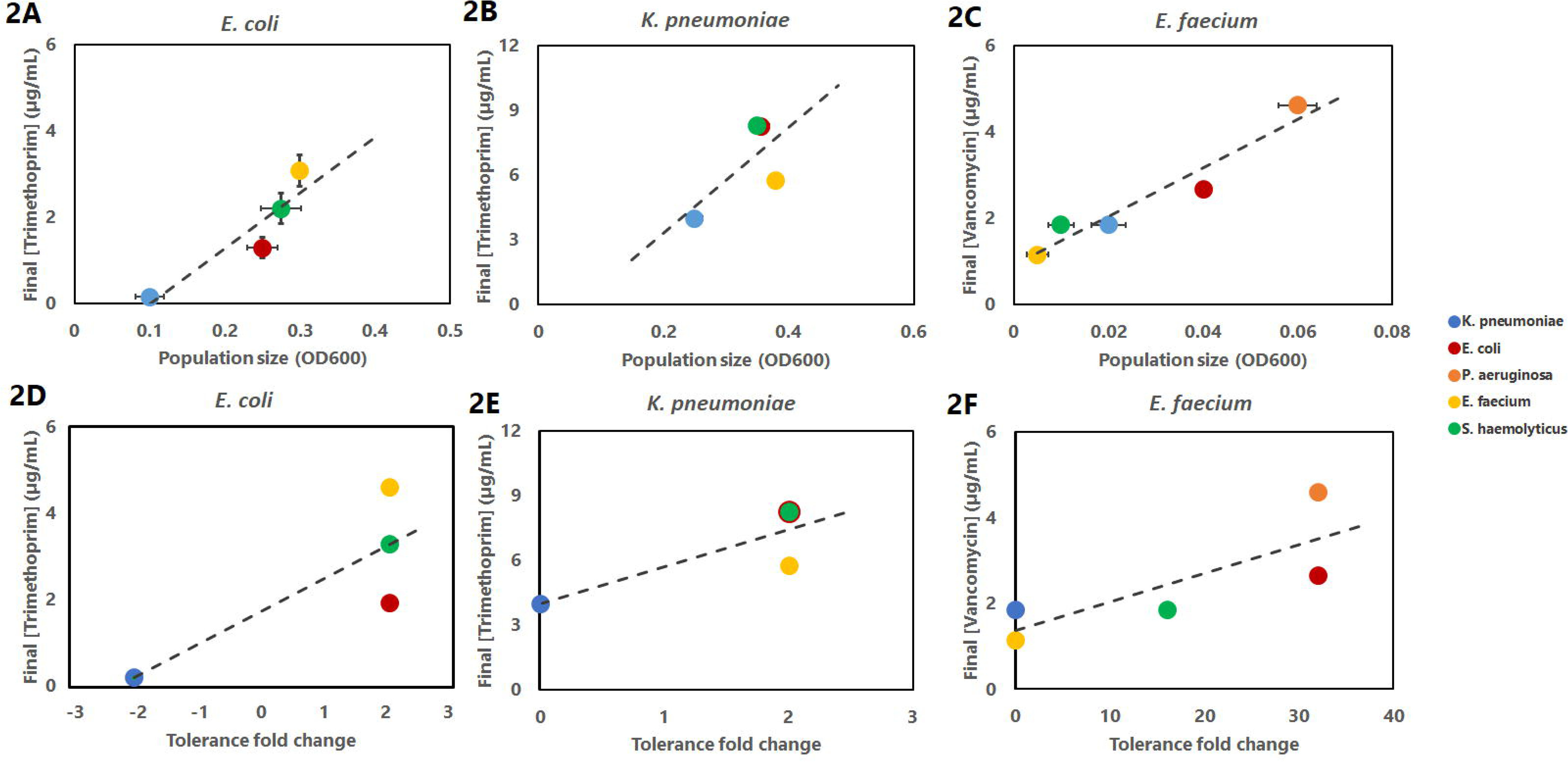
Correlation of antibiotic concentration at the final transfer with ecological interactions; WT population size in conditioned medium, and the fold change of antibiotic tolerance in conditioned medium. Colors indicate the conditioned medium in which the focal species were cultured, with blue indicating *K. pneumoniae* conditioned medium, red *E. coli* conditioned medium, yellow *E. faecium* conditioned medium and green *S. haemolyticus* conditioned medium. The *E. faecium* mutants were also evolved in *P. aeruginosa* conditioned medium, represented by orange markers. **A**, Trimethoprim concentration (ratio sulfamethoxazole:trimethoprim is 5:1) at final transfer of *E. coli* lineages in the serial propagation experiment correlated with the population size interactions of the wild type in the absence of antibiotics (*R^2^*=0.86, *N=*20, *P<*0.001, means ± s.e.m.). **B**, Trimethoprim concentration (ratio sulfamethoxazole:trimethoprim is 5:1) at final transfer of *K. pneumoniae* lineages in the serial propagation experiment correlated with the population size interactions of the wild type in the absence of antibiotics (*R^2^*=0.45, *N=*20, *P*=0.017, means ± s.e.m.). **C**, Vancomycin concentration at final transfer of *E. faecium* lineages in the serial propagation experiment correlated with the population size of the wild type in the absence of antibiotics (*R^2^*=0.93, *N=*25, *P*<0.001, means ± s.e.m.). **D**, Correlation between antibiotic concentration at final transfer of *E. coli* mutants of in the serial propagation experiment versus the fold change of the antibiotic tolerance of the wild-type *E. coli* in the conditioned media compared to artificial urine medium (*R^2^*=0.50, *N=*20, *P*<0.001). **E**, Correlation between antibiotic concentration at final transfer of *K. pneumoniae* mutants of in the serial propagation experiment versus the fold change of the antibiotic tolerance of the wild-type *K. pneumoniae* in the conditioned media compared to artificial urine medium (*R^2^*=0.68, *N=*20, *P*<0.001). **F**, Correlation between antibiotic concentration at final transfer of *E. faecium* mutants of in the serial propagation experiment versus the fold change of the antibiotic tolerance of the wild-type *E. faecium* in the conditioned media compared to artificial urine medium (*R^2^*=0.64, *N=*25, *P*<0.001).

Because phylogenetically similar species are expected to have the most niche overlap, we hypothesize that the conditioned medium of phylogenetically similar species leads to the most niche overlap, and consequently, the smallest population sizes, and hence the slowest evolution, as established above. Indeed, we find that the rate and level of antibiotic resistance evolution is slowest in the conditioned medium of the focal isolate for two of the three focal species (*K. pneumoniae* in *K. pneumoniae* conditioned medium, and *E. faecium* in *E. faecium* conditioned medium, Figure 1B,C). Moreover, for two of the three focal isolates the rate of evolution is fastest in the conditioned medium of the isolates that are phylogenetically least related (Supplementary Figure 1), *E. faecium* evolves fastest in the Gram-negative conditioned media (*H*=16.39, *P*<0.005, Kruskal-Wallis), and *E. coli* evolves fastest in the Gram-positive conditioned media (*H*=12.09, *P*<0.005). For *K. pneumoniae* we observe no such niche effects (*H*=0.006, *P*=0.94), other than the slowest evolution in its own conditioned medium. Alternatively, the conditioned medium of donor species may affect population sizes by other means than niche overlap. For instance, we find that *K. pneumoniae* conditioned medium was associated with the highest pH (pH 8), compared to the other conditioned media (pH 7.5 *S. haemolyticus and P. aeruginosa;* pH 7.25 *E. faecium* and *E. coli* conditioned medium). This increased alkaline condition, was associated with the slower evolution of *E. coli* in *K. pneumoniae* conditioned medium (40), but not the slower evolution of *E. faecium*, which is known to tolerate a wider range of pH conditions (Figure 1A,C) (41,42). This suggests that niche specific effects such as avoidance of competition, but also more general niche effectors, such as pH, could affect bacterial interactions that promote the divergence of antibiotic resistance evolution, as also observed in other studies (43–46).

We previously found that growth-mediating effects of bacterial interactions in the absence of antibiotics correlate with the level of antibiotic tolerance (6). Here we define the tolerance as the level of an antibiotic that a population can sustain under the influence of bacterial interactions (6). The fold change in antibiotic tolerance was calculated based on the change in growth inhibiting antibiotic concentration of conditioned media compared to unconditioned medium. Here, we find that the concentration of antibiotics at the last transfer of the experiment correlates positively with the antibiotic tolerance levels conferred by the conditioned media (*R^2^*=0.50, 0.68 and 0.64 for *E. coli* (*N*=20), *K. pneumoniae* (*N*=20), and *E. faecium* (*N*=25) resp. for all *P*<0.001, Figure 2D-F). Possibly, these effects of conditioned media on antibiotic tolerance are driven by affecting the population size, similar to the inoculum effect in assays measuring the minimal inhibitory concentration of certain antibiotics (11,47–49).

### Ecological interactions do not affect the number of genetic changes

To investigate the consequences of these ecological interactions on evolutionary trajectories, we determined the genetic changes in these evolved lineages through the sequencing of three clones per evolved lineage (Materials and Methods). A fundamental determinant of evolutionary trajectories is the population size, as it affects both the supply of mutations and the efficacy of natural selection to select beneficial mutations (17,20,50). Therefore, effects on population size may lead to different mutations being selected (51,52).

To quantify the impact of natural selection in our populations, we measured the dN/dS ratio (35,53) (Materials and Methods). Generally, we find dN/dS ratios >> 1, resp. *E. coli* 4.5, *K. pneumoniae* 7.2, *E. faecium* 17, indicating strong positive selection in all populations (chi-squared test to test deviation from expected dN/dS in the absence of selection, *P*=0.024, 0.004, 0.001, for respectively *E. coli*, *K. pneumoniae*, *E. faecium*, Supplementary Table 1, Materials and Methods).

SNPs outnumbered other mutations in the evolved isolates (Supplementary Figure 2A-C). Yet, we find no positive correlation between the number of SNPs in the end-point mutants and population size for *E. coli* and *E. faecium* (*R^2^*=0.18 and 0.47, *P*=0.17, 0.20, resp., Supplementary Figure 3A,C), and a negative correlation for *K. pneumoniae* (*R^2^*=0.70, *P*<0.001, Supplementary Figure 3B). Other genetic differences were also observed, such as deletions, inversions and copy-number differences. Moreover, in several of these *K. pneumoniae* lineages one of the plasmids was entirely lost. Especially for the second plasmid (Supplementary Figure 4B, Supplementary Table 2), we observed that blocks of the plasmid were deleted during evolution. These mutational patterns were not observed to be associated with specific conditioned media.

We find no positive correlation between the different conditioned media associated with different population sizes, and the total number of genetic changes (*R^2^*=0.023 and 0.03, 0.15, *P*=0.68, 0.59, 0.19 for *E. coli, K. pneumoniae* and *E. faecium* lineages respectively, Supplementary Figure 5A-C). Therefore, population size effects did not affect the number of genetic changes in the end-point mutants.

### Ecological interactions do affect the types of genetic changes

To test whether different conditioned media lead to the incorporation of different genetic changes in the adaptive trajectories of the three focal species, we compared the genetic similarity of isolates evolved in the same with those evolved in different conditioned media. We calculated the *H-*index (17), as a measure for the fraction of mutated genes that two evolved isolates have in common (Materials and Methods). If we calculate this index at the level of Clusters of Orthologous Groups (COGs) (54), i.e. genes grouped by function, we find more parallelism among replicate populations evolved within the same conditioned media compared to those same lineages in different conditioned media (*P*=0.043, 0.043, 0.032 (Welch’s t-test), excluding self-comparisons, for *E. coli*, *K. pneumoniae* and *E. faecium,* respectively, Figure 3). This indicates that the parallel evolution observed (Figure 1), compliments the parallelism observed at the functional level in the same conditioned media.

**Figure 3.**
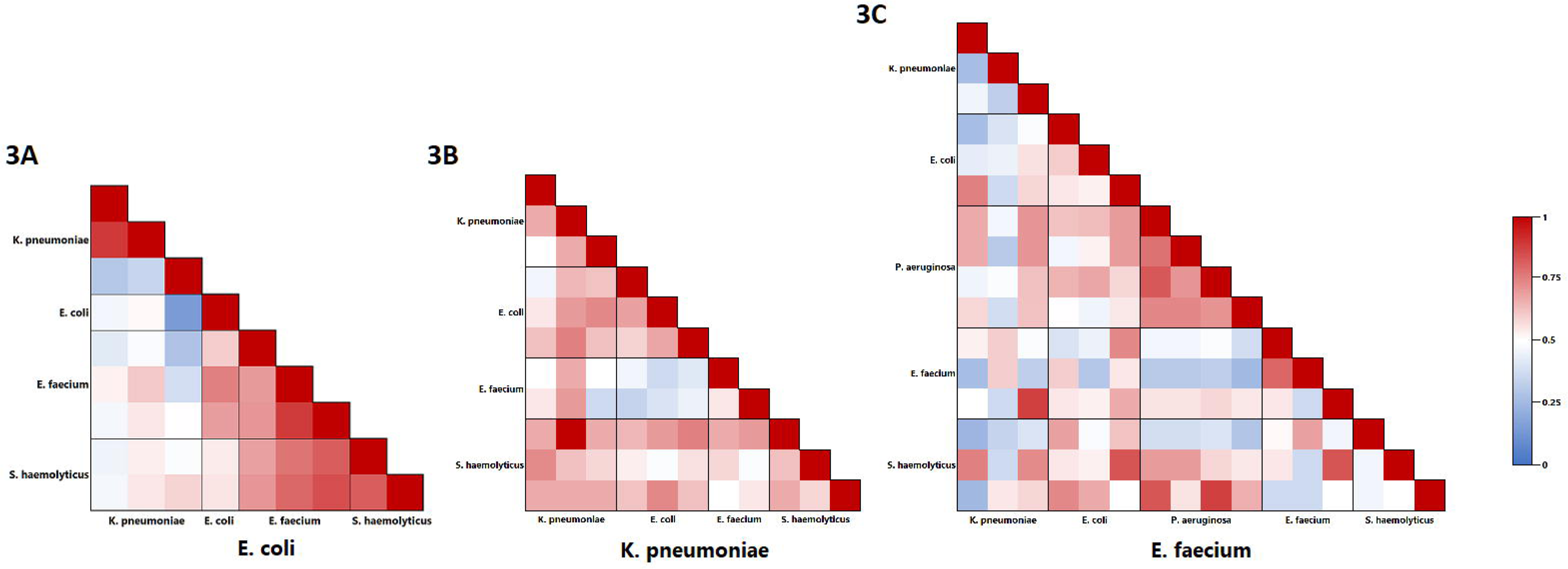
Clustering and mutation repeatability of the final propagation step mutants clustered on COG-level. Higher values indicate a higher parallelism of the two genotypes compared. Three mutant lineages were sequenced per conditioned medium (four for *E. faecium* mutants evolved in *P. aeruginosa* conditioned medium). Comparisons in the blocks around the diagonal in each figure show the intra-conditioned medium comparisons, whereas the other blocks show the inter-conditioned media comparisons. **A,** *H*-index of mutation repeatability at COG-level of *E. coli* evolved in the four conditioned media. Mutator lineages are not taken into account (Supplementary Figure 1A). **B,** *H*-index of mutation repeatability at COG-level of *K. pneumoniae* evolved in the four conditioned media. Mutator lineages are not taken into account (Supplementary Figure 1B). **C,** *H*-index of mutation repeatability at COG-level of *E. faecium* evolved in the five conditioned media.

For *E. coli*, particularly the lineages that evolved in the Gram-positive conditioned media of *S. haemolyticus* and *E. faecium* display a high degree of parallelism at the COG-level (*H*>0.6, Figure 3A, *P*<0.05 compared to the other *H*-values, Welch’s t-test). This pattern is conserved at the gene-level for lineages evolved in *S. haemolyticus* conditioned medium (Supplementary Figure 6A). The high genetic parallelism corresponds to a similar phenotypic growth response of *E. coli* replicates to these conditioned media (Figure 2D), and their partially overlapping evolutionary trajectories (Figure 1A). Interestingly, the population size of *E. coli* in the conditioned medium of itself is also similar to the population size in the conditioned medium of *S. haemolyticus* (Figure 2A, *P*>0.05 when comparing the two set, Welch’s t-test). Yet, the *H-*index values that correspond to the *E. coli* lineages evolved in these Gram-positive (*S. haemolyticus)* and Gram-negative (*E. coli)* conditioned media are relatively low (*H*<0.6), indicating lower parallelism within these conditions. This shows that population size alone cannot explain the level of parallelism, which suggests that the constituents of the conditioned medium, also play a role in the genetic divergence of these evolutionary trajectories.

For *E. faecium*, we find a particularly strong parallelism at the COG-level for evolved lineages in *P. aeruginosa* conditioned medium, but less so for other conditioned media (Figure 3C). At the gene-level, this strong parallelism for evolved lineages in *P. aeruginosa* conditioned medium is not present (Supplementary Figure 6C). The high level of parallelism among *E. faecium* lineages evolved in *P. aeruginosa* conditioned medium coincides with the largest population sizes, and the highest rate of evolution (Figure 1C). Yet, the level of parallelism of *E. faecium* evolved in the conditioned media of *K. pneumoniae* and *S. haemolyticus* is lower, despite similar resistance trajectories and population sizes (*H*-index means of 0.75 for *P. aeruginosa,* and 0.35 (*P*=0.022) and 0.48 (*P*=0.013) for *K. pneumoniae* and *S. haemolyticus* resp., Welch’s t-tests *H*-index comparison of *E. faecium* in the respective conditioned media) (Figure 1B, 3B). This again suggests that particular metabolic constituents of the conditioned medium also play a role in the genetic divergence.

For *K. pneumoniae*, we find consistently high *H*-indices (*H*>0.6) at the COG-level across all conditions, but a little lower index for the comparison of the isolates evolved in *E. faecium* and *E. coli* conditioned media (Figure 3B). In addition, at the gene-level we also find relatively high parallelism across all conditions (although slightly lower than at the COG-level), especially for the isolates evolved within *S. haemolyticus* and *K. pneumoniae* conditioned medium (Figure 3B). These findings highlight the rather limited effect of conditioned media on the antibiotic resistance evolution of *K. pneumoniae* (Figure 2B).

As observed before (17,55), the *H*-indices show a higher parallelism at the COG-level than at the gene-level (Figure 3, Supplementary Figure 6). This suggests that there is genetic redundancy in the functionalities that are selected under these particular ecological circumstances. Generally, the higher *H*-index of lineages evolved within the same conditioned medium (along the diagonal axis in Figure 3) shows that the antibiotic resistance evolution in the conditioned media leads to parallel genotypic (Figure 3) and phenotypic (Figure 1) adaptation under the influence of species-specific bacterial metabolites.

### Conditioned medium causes specific genetic changes

Zooming in on the genetic changes, we investigated whether the species-specific ecological interactions also lead to the selection of different resistance mechanisms. We do find that in particular conditioned media specific mutations occur repeatedly, whereas others do not (Figure 4A-C). In accordance with other research, we hypothesized that in larger populations, larger effect-size mutations are selected, because these are not limited in their mutation supply and are under strong selection, whereas in smaller populations, different types of smaller effect size mutations would be selected (17,19). Indeed, we find that the lineages with the smallest population size, which is *E. faecium* cultured in its own conditioned medium, repeatedly lacked mutations in known antibiotic resistance targets (Figure 4C) (56–58). When evolved in other conditioned media, with larger population sizes, they reached higher levels of resistance and incorporated mutations in well-known vancomycin resistance targets, *VanR, VanS* (56,59,60), or both. *VanS* is a sensor histidine kinase that detects vancomycin and activates *VanR* (56,59,61). *VanR* is a transcriptional activator, regulating the transcription of the vancomycin-resistance operon together with regulating transcription of the vanHAX operon, resulting in higher tolerance due to changes in peptidoglycan formation (56,60). Indeed, lineages that had mutations in well-known resistance targets were generally associated with larger population sizes and higher resistance levels (Figure 4C, 2C).

**Figure 4.**
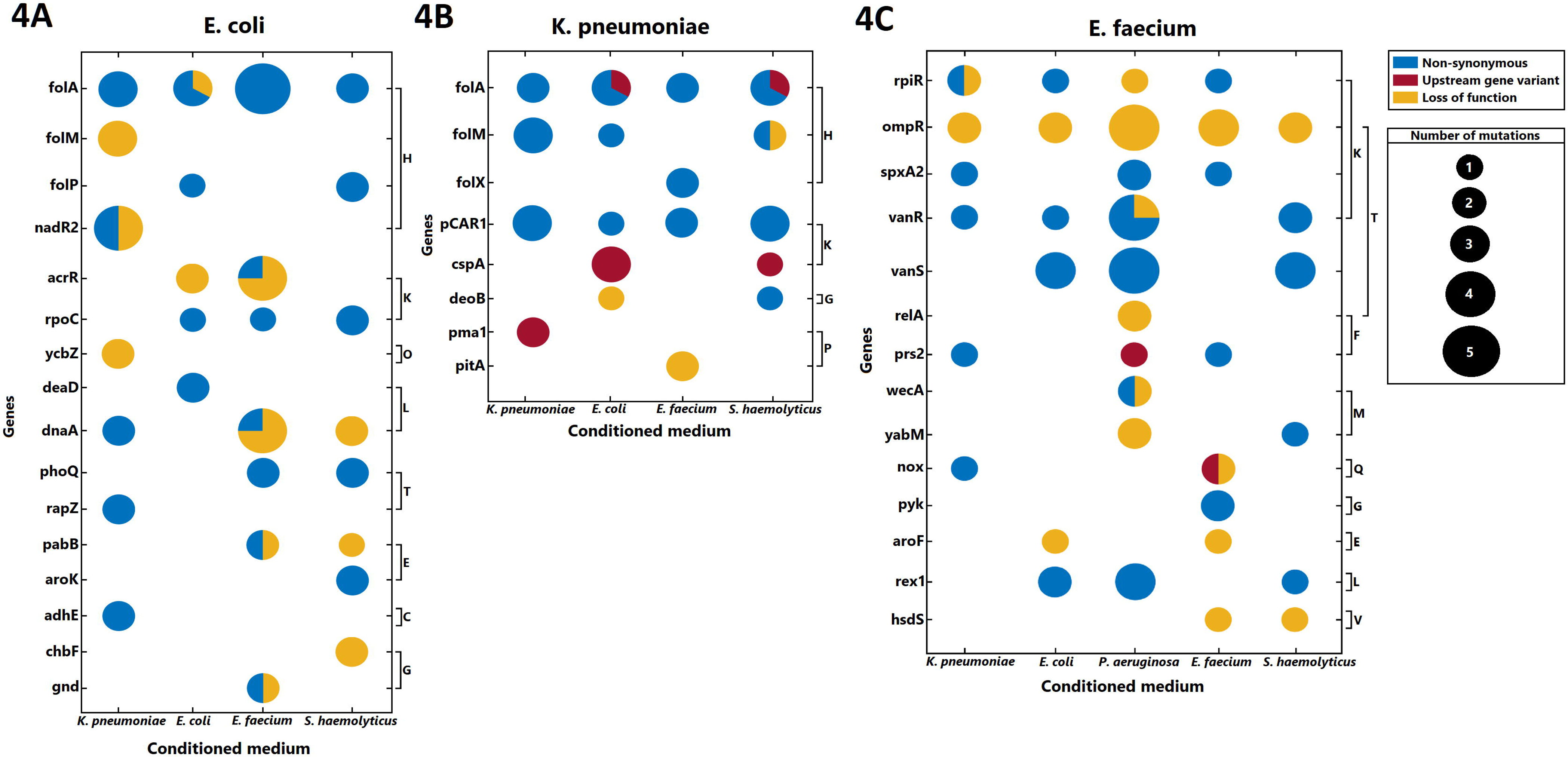
Evolved genetic targets separated by type of mutation. Genetic targets shown were hit multiple times (at least twice) in either replicate populations or environment, or both. Size of the spheres indicate the number of unique mutations in the specific gene that were observed. Genes are separated into their respective COG functional group as shown on the right side of the figures, with each letter corresponding to a different COG-group. COG-groups: C – Energy production and conversion, E – Amino acid transport and metabolism, F – Nucleotide transport and metabolism, G – Carbohydrate transport and metabolism, H – Coenzyme transport and metabolism, K – Transcription, L – Replication, recombination and repair, M – Cell wall/membrane/envelope biogenesis, O – Posttranslational modification, protein turnover, chaperones, P – Inorganic ion transport and metabolism, Q – Secondary metabolites biosynthesis, transport and catabolism, R – General function prediction only, T Signal transduction mechanisms, and V – Defense mechanisms. **A,** Genetic targets in *E. coli* evolved in four different conditioned media. See Supplementary Table 3 for a list of the genes and proteins. **B,** Genetic targets in *K. pneumoniae* evolved in four different conditioned media. See Supplementary Table 4 for a list of the genes and proteins. **C,** Genetic targets in *E. faecium* evolved in five different conditioned media. See Supplementary Table 5 for a list of the genes and proteins.

The two Gram-negative lineages (*K. pneumoniae* and *E. coli*) were selected for resistance to trimethoprim-sulfamethoxazole. All sequenced isolates had mutations in *folA*, dihydrofolate reductase (Figure 4A,B, Supplementary table 3-4), which is involved in the folate pathway that is essential for DNA synthesis and is inhibited by trimethoprim-sulfamethoxazole (62). *FolA* is the direct target of trimethoprim. One lineage of *E. coli* evolved in *E. coli* conditioned medium even had a 20-fold amplification of the copy-number of this gene.

Both *K. pneumoniae* and *E. coli* incorporated *folM* mutations. *FolM* encodes an isozyme of dihydrofolate reductase and is also involved in DNA synthesis. Inhibition of this gene causes the upregulation of *folA,* but usually also requires resistance mutations in *folA* to be effective (63). *E. coli* incorporated *folM* mutations only in lineages that evolved slowly in the *K. pneumoniae* conditioned medium (Figure 4A). Yet, in *K. pneumoniae*, *folM* mutations appeared in the slowest evolving lineages in *K. pneumoniae* medium, but also in the fastest, namely the *E. coli* and *S. haemolyticus* conditioned media.

In *K. pneumoniae* in *E. faecium* conditioned medium, *folX* was targeted instead of *folM* (Figure 4B). *E. coli* lineages evolved in *E. coli* and *S. haemolyticus* conditioned medium incorporated mutations in yet another gene in the folate pathway, *folP* (64) (Figure 4A). All these mutations in the folate pathway correspond to clinically relevant mutations (Supplementary text 2). The observation that different genes within the folate pathway are targeted in the different conditioned media suggests that ecological interactions may cause different trajectories of antibiotic resistance evolution.

### Antibiotic resistance and cost of resistance depend on the genetic background and the ecological interactions

To assess whether the lineages evolved to the conditioned media or to the antibiotics, we assessed the growth of the end-point mutants of evolved lineages in each conditioned medium, and measured the antibiotic resistance level in the absence of conditioned media. The relationship between the evolved the minimal inhibitory concentration (MIC) and the antibiotic concentration at the final transfer shows a negative correlation between the MIC and the end-point antibiotic concentrations for *K. pneumoniae*, and a positive correlation for *E. coli* and *E. faecium* (Supplementary Figure 7). For these latter focal isolates the highest resistance evolved in the most and fastest evolving populations (Supplementary Figure 7A,C), whereas for *K. pneumoniae* resistance evolved fast in all conditioned media (Figure 1), but the highest resistance evolved during the slowest adaptation (Supplementary figure 7). We also find a cost of resistance of most mutants, as the growth is mostly below that of the ancestor (Supplementary figure 8). We find no correlation between the MIC in the absence of conditioned medium, and the growth of the mutants.

Adaptation to the conditioned medium is shown by the fact that the growth of mutants evolved in a particular conditioned medium mostly (9 out of 13) lie in the upper right quadrant (Supplementary Figure 9-11, for those 9 *P<0.05*, one-way ANOVA), showing improved growth in those conditioned media, compared to the growth in other conditioned media. The fact that resistance evolution is subject to ecological interactions, is further exemplified by the growth of end-point clones of *E. coli* evolved in *E. faecium* conditioned medium (Supplementary Figure 9C,E), in the presence and absence of antibiotics. This shows that costs and benefits of accrued mutations in these lineages are dependent on both environmental factors, the presence of antibiotics and the conditioned medium. Evolution of antibiotic resistance in the presence of microbial interactions is therefore a truly eco-evolutionary process.

## Discussion and conclusion

We find that the bacterial interactions mediated by conditioned medium differentially altered the rate and mutational trajectories of antibiotic resistance evolution of uropathogenic isolates *E. coli*, *K. pneumoniae* and *E. faecium*. These results are in line with other work that showed that community composition can alter the selection for antibiotic resistance, for instance through modulating population densities via competitive or cooperative interactions (49,65–70). In the conditioned media environment, cultured bacteria were not in physical contact with the bacteria used to create conditioned media. Such contact-mediated interactions may affect eco-evolutionary outcomes, since two populations of bacteria can respond to the presence and the produced metabolites of the other when in the same environment (71,72). Moreover, higher-order interactions between more than two bacterial species could potentially affect growth (73,74). Lastly, the donor species were not cultured in the presence of antibiotics, these could therefore not ‘soak’ (75) or degrade the level of antibiotics (76). Future studies should address such additional factors that can affect the evolution of antibiotic resistance.

Overall, we found parallelism in lineages evolved under the same ecological conditions (Figure 1), and that resistance mutations and other functional targets selected were dependent on these conditions (Figure 3, 4). Initial standing variation cannot account for this pattern, since different mutations in the same genes were incorporated (Supplementary tables 3-5). Parallelism has been observed to depend on environmental conditions before, such as resource availability (77), ecological interactions (78) and population size of evolving lineages (17), but the phenotypic parallelism does not always correspond to molecular parallelism (79). This work shows that ecological interactions, which may be through avoidance of resource competition (utilizing different niches), cross-feeding, or pH changes, and potentially other antibiotic tolerance effects (6,45,66), can drive both phenotypic and genetic parallel antibiotic resistance evolution.

*K. pneumoniae* showed the least variation across conditioned media, in terms of growth, tolerance as well as the rate and final level of adaptation. It also showed the highest parallelism within each conditioned medium at the genetic level, which correlated with the overall consistently large population sizes and therefore a smaller effect on mutation supply (19,20). In contrast, *E. coli* and *E. faecium* showed more variation across conditioned media. Particularly, *E. faecium* lineages that evolved in the smallest population sizes (in *E. faecium* conditioned medium), did not incorporate such large effect resistance mutations. These lineages also showed the highest conditioned medium dependence at the genetic level, which was not always associated with the population size effects. This suggests that for these species different conditioned media may have a differential effect on the functions under selection, such that the underlying shape of the adaptive landscapes of resistance evolution would be more environmentally dependent (80,81), or that the mechanisms that confer evolutionary benefits are different depending on the antibiotic concentration (82). Following the mutational trajectories during the emergence of antibiotic resistance evolution could lead to more insight in the ecological impact on resistance by identifying genotypes that alter these trajectories (83–87).

Concluding, antimicrobial resistance is turning into a “silent pandemic” (58), as infections caused by resistant microorganisms fail to respond to conventional treatment, resulting in prolonged illness, greater risk of death and higher healthcare costs (88). Ecological interactions, mediating population sizes and other ecological factors, can explain the deviating antibiotic resistance levels as measured in isolation (e.g., in the clinic) (6,47,69). We find that such interactions between bacteria, and conferred antibiotic tolerance, affect the mode, the genetic functions selected, and the tempo, the rate of antibiotic resistance evolution. The identification of the nature of such ecological interactions can aid in forecasting the evolutionary potential of antibiotic resistance evolution. Further knowledge of such predictive parameters can improve decision making, which will be crucial for developing interventions, and for curbing the emergence and further evolution of antibiotic resistance (89).

## Acknowledgements

We thank Ineke Heikamp-de Jong for help with the DNA extraction, and Bart Thomma for the use of the MLII laboratory at Wageningen University. MGJDV was supported by an NWO VENI Fellowship (863.14.015), ZonMw ETH grant (40-43500-98-4049/435004009) and L’Oreal UNESCO LNVH NIAS-KNAW For Women in Science Fellowship, part of this work was performed at NIAS-KNAW. LEZ was supported by the Adaptive Life program of the Faculty of Science and Engineering, within GELIFES.

## Contributions

Conceived and designed the study: LEZ JAGMDV BJZ MGJDV. Performed the experiments: LEZ ADF MGJDV. Analyzed the data: LEZ JVDH MGJDV. Wrote the manuscript: LEZ JVDH ADF JAGMDV MGJDV.

## Data availability statement

Bacterial sequence data have been deposited with links to BioProject accession number PRJNA1103400 in the NCBI BioProject database. The other datasets generated during and/or analyzed during the current study are available from the corresponding author on reasonable request.

## Notes

http://www.ncbi.nlm.nih.gov/bioproject/1103400

### Competing Interest Statement

The authors have declared no competing interest.

### Summary of Updates

Figure 1 revised, added maximum rate of evolution; Discussion revised and updated; Figure 'Amount and types of mutations in end-point mutants' moved to the supplementary information; Added section 'Antibiotic resistance and cost of resistance depend on the genetic background and the ecological interactions'; Supplementary files updated, including extended supplementary materials and methods; Added phylogenetic tree of 16S sequences of isolates to supplementary information;

