## Supplementary Information for "Microbial interactions affect the tempo and mode of antibiotic resistance evolution"

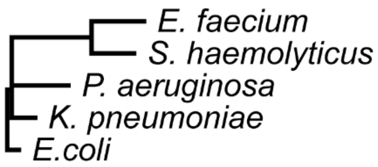

**Supplementary Figure 1 | Phylogenetic relatedness of isolates used in this study, based on 16S rDNA.**

Two clusters can be observed, one cluster consisting of the Gram-positive species *E. faecium* and *S. haemolyticus,* and the other consisting of the Gram-negative species *P. aeruginosa*, *K. pneumoniae* and *E. coli* (Materials and Methods). For *E. coli* the conditioned medium generated from Gram-positive species have a more positive effect on the evolution ($H$=12.09, $P$<0.005). For *E. faecium* the Gram-negative species have a more positive effect on the evolution ($H$=16.39, $P$<0.005). This is not the case for *K. pneumoniae* ($H$=0.006, $P$=0.94). Significance was calculated by Kruskal-Wallis rank sum test.

S2A

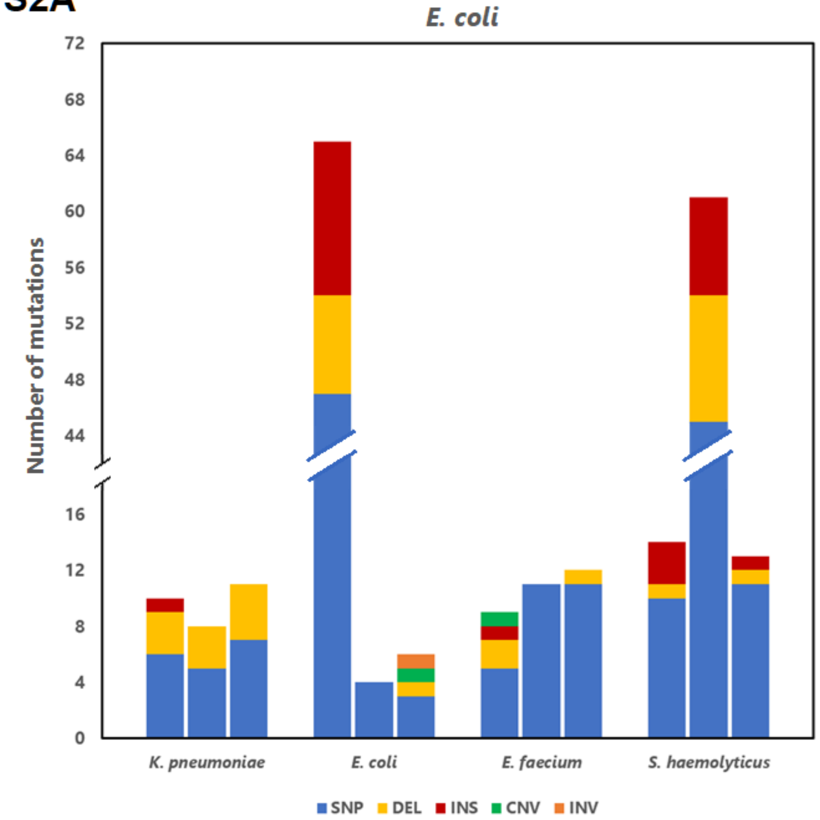
E. coli
Number of mutations
0
4
8
12
16
44
48
52
56
60
64
68
72
K. pneumoniae
E. coli
E. faecium
S. haemolyticus
SNP
DEL
INS
CNV
INV

S2B

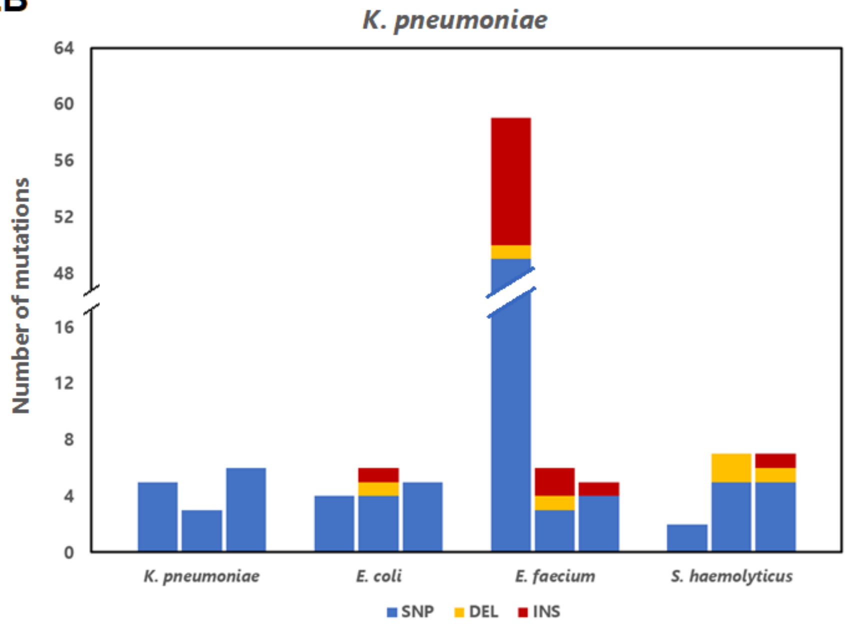
K. pneumoniae
Number of mutations
0
4
8
12
16
48
52
56
60
64
K. pneumoniae
E. coli
E. faecium
S. haemolyticus
SNP
DEL
INS

S3B

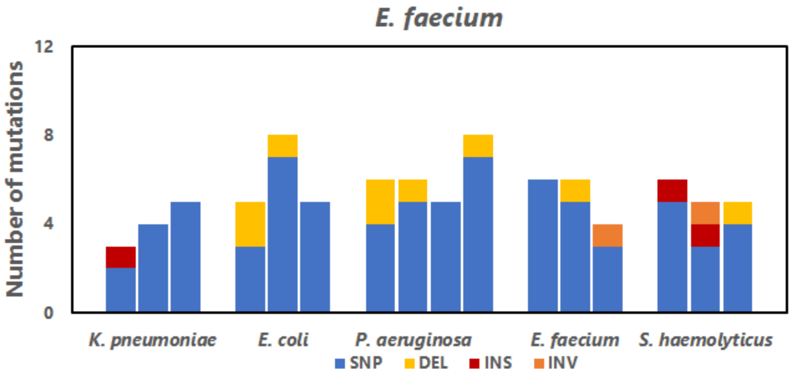
E. faecium
Number of mutations
0
4
8
12
K. pneumoniae
E. coli
P. aeruginosa
E. faecium
S. haemolyticus
SNP
DEL
INS
INV

**Supplementary Figure 2 | Amount and types of mutations in end-point mutants. A,** All mutations in the *E. coli* mutants evolved in the conditioned media shown on the x-axis per replicate lineage. Two mutator strains were identified, shown here by the broken y-axis/bars. **B,** All mutations in the *K. pneumoniae* mutants evolved in the conditioned media shown on the x-axis per replicate lineage. One mutator strain was identified, shown here by the broken y-axis/bars. **C,** All mutations in the *E. faecium* mutants evolved in the conditioned media shown on the x-axis per replicate lineage. The three hyper-mutator lineages accumulated many more mutations compared to the other evolved lineages of these species: around 60 mutations in comparison to the average of ≈10, for *E. coli* and *K. pneumoniae* (Supplementary Figure 3D, E, Poisson-test, $P < 0.001$). For this reason, these lineages were removed from the SNP count analysis. The sequences of the *E. coli* hyper-mutator lineages showed large deletions (>3KB) in the region which includes *mutH*, a gene involved in mismatch repair (1), which points to a role for *mutH* in the generation of these hyper mutator lineages. The *K. pneumoniae* hyper-mutator lineage appeared in *E. faecium* conditioned medium, was associated with a missense SNP mutation in *mutS*. *K. pneumoniae* (B) and *E. faecium* (C) are evolved under different circumstances (e.g., different intrinsic population sizes, ~$10^8$ and ~ $10^4$ respectively), yet differences in intrinsic mutation rates cannot explain their similar number of mutations (t-test, $N = 6$, $P = 0.13$, Supplementary Table 3, Materials and Methods).

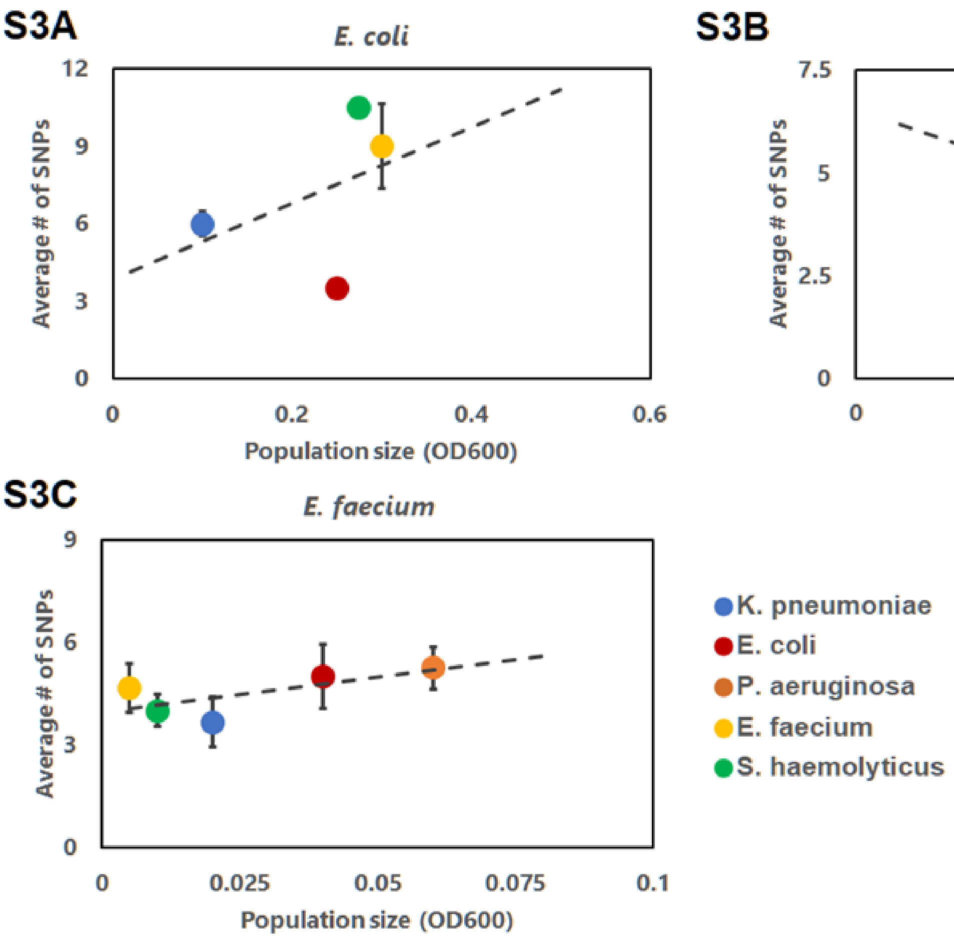

**Supplementary Figure 3 | Mutation supply, number of SNPs and type of mutations in final propagation step mutants.** Colors indicate the conditioned medium in which the focal species were cultured, with blue showing mutants evolved in *K. pneumoniae* conditioned medium, red showing mutants evolved in *E. coli* conditioned medium, yellow showing mutants evolved in *E. faecium* conditioned medium, and green showing mutants evolved in *S. haemolyticus* conditioned medium. For *E. faecium,* mutants were also evolved in *P. aeruginosa* conditioned medium, which is shown with the orange marker. The error bars show the standard error of the mean for the number of SNPs in the non-mutator trains. **A,** The average number of single nucleotide polymorphisms (SNPs) in the clones of evolved *E. coli* in the conditioned media correlated with the population size of the wild type in those conditioned media ($R^2$=0.18, $N$=12, $P$=0.17, means ± s.e.m.). **B,** The average number of SNPs in the clones of *K. pneumoniae* in the conditioned media correlated with the population size of the wild type in those conditioned media ($R^2$=0.704 $N$=12, $P$<0.001, means ± s.e.m.). **C,** The average number of SNPs in the clones of evolved *E. faecium* in the conditioned media correlated with the population size of the wild type in those conditioned media ($R^2$=0.76, $N$=16, $P$=0.1989, means ± s.e.m.).

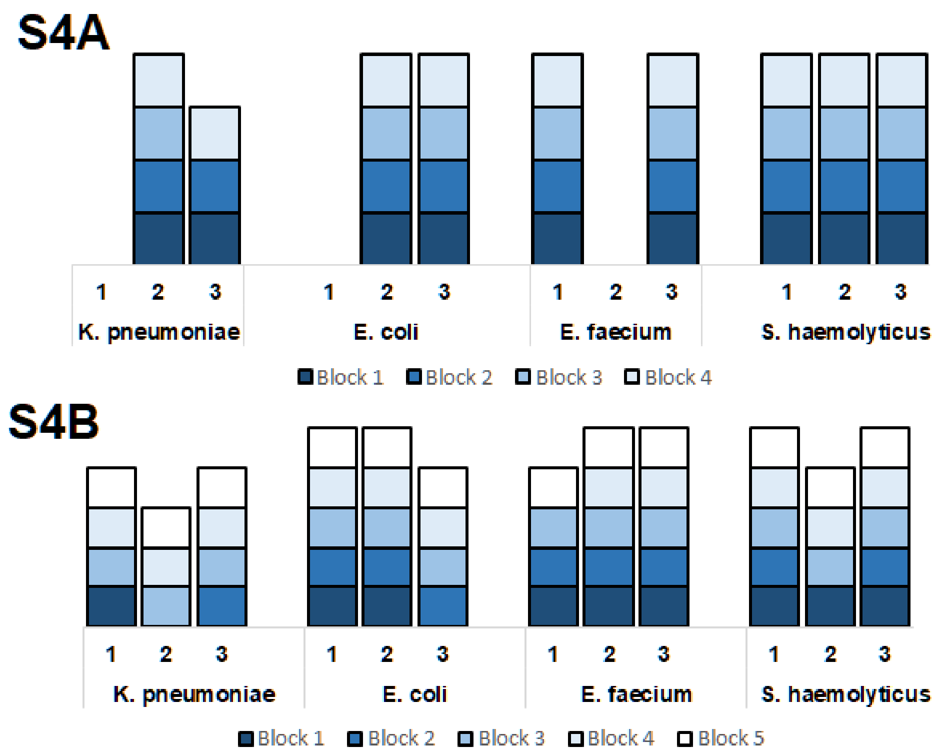

**Supplementary Figure 4 | Plasmid changes in *K. pneumoniae.*** Changes in the presence of gene blocks on the **A** first plasmid and **B** second plasmid in *K. pneumoniae* mutants. Three replicates per conditioned medium background are shown on the x-axis. Genes present on these blocks can be found in Supplementary Table 1. These two plasmids that had copy-number differences in several evolved lineages. Parts of these two plasmids are identical to sequences found in Genbank, with part of the first plasmid being identical to a *K. pneumoniae* plasmid (Isolate KSB1_10B-sc-2280232, accession no. LR890699) and part of the second plasmid being identical to a plasmid in a *K. oxytoca* (Isolate RHBSTW-00493, accession no. CP056454)(2).

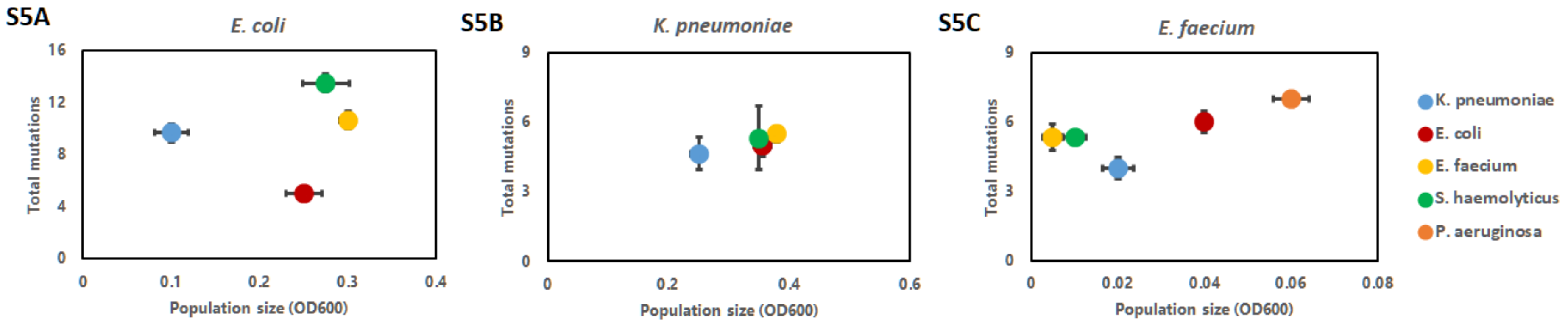

**Supplementary Figure 5 | Correlation of population size interactions with the total number of mutations.** Population sizes of the wild types were measured in the conditioned media, as indicated by the different colors, with data showing means ± s.e.m. **A,** *E. coli* replicates show no correlation with $R^2$=0.023 and $P$ = 0.68, $N$=10. **B,** *K. pneumoniae* replicates showed no correlation with $R^2$=0.03 and $P$=0.59, $N$=11. **C,** *E. faecium* replicates show a small but insignificant positive correlation between the population size interactions and mutations with $R^2$=0.15 and $P$=0.19, $N$=13.

S6A

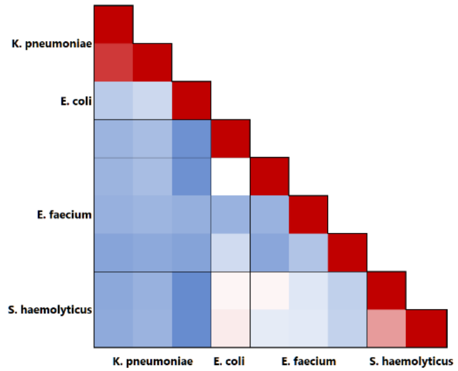
K. pneumoniae
E. coli
E. faecium
S. haemolyticus
K. pneumoniae
E. coli
E. faecium
S. haemolyticus

S6B

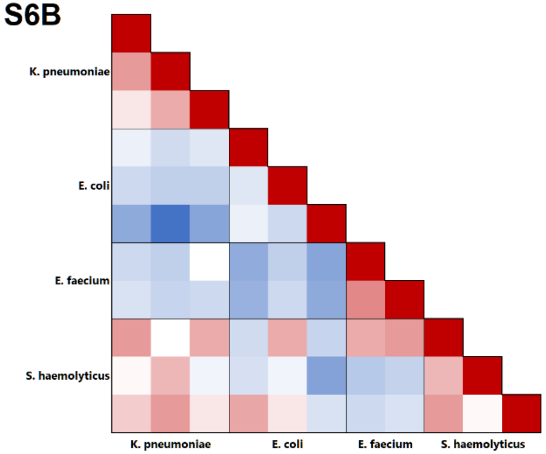
K. pneumoniae
E. coli
E. faecium
S. haemolyticus
K. pneumoniae
E. coli
E. faecium
S. haemolyticus

S6C

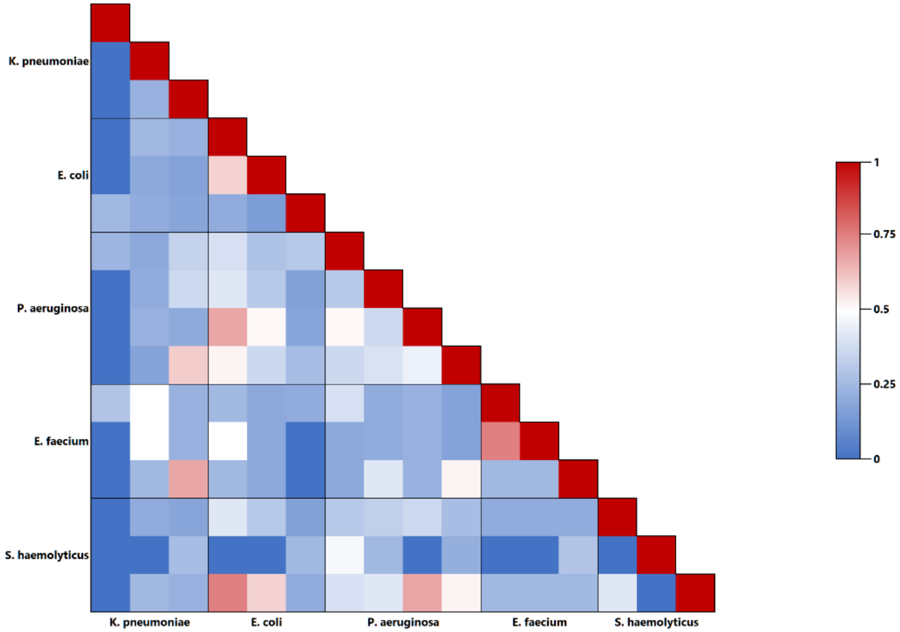
K. pneumoniae
E. coli
P. aeruginosa
E. faecium
S. haemolyticus
K. pneumoniae
E. coli
P. aeruginosa
E. faecium
S. haemolyticus
1
0.75
0.5
0.25
0

**Supplementary Figure 6 | Clustering and mutation repeatability of the final propagation step mutants on gene-level.** Higher values indicate a higher parallelism of the two genotypes compared. Comparisons in the blocks around the diagonal in each figure show the intra-conditioned medium comparisons, while the other blocks show the inter-conditioned media comparisons. **A,** *H*-index of mutation repeatability at gene-level of *E. coli* evolved in the four conditioned media. Mutator lineages are not taken into account (Figure S1A). **B,** *H*-index of mutation repeatability at gene-level of *K. pneumoniae* evolved in the four conditioned media. Mutator lineages are not taken into account (Figure S1B). **C,** *H*-index of mutation repeatability at gene-level of *E. faecium* evolved in the five conditioned media. Comparisons of *H*-indices within conditioned media was not significantly different from between conditioned media for all three focal species (*P*=0.4334, 0.4341, 0.3242 for *E. coli*, *K. pneumoniae* and *E. faecium* respectively (*Welch's t-tests*).

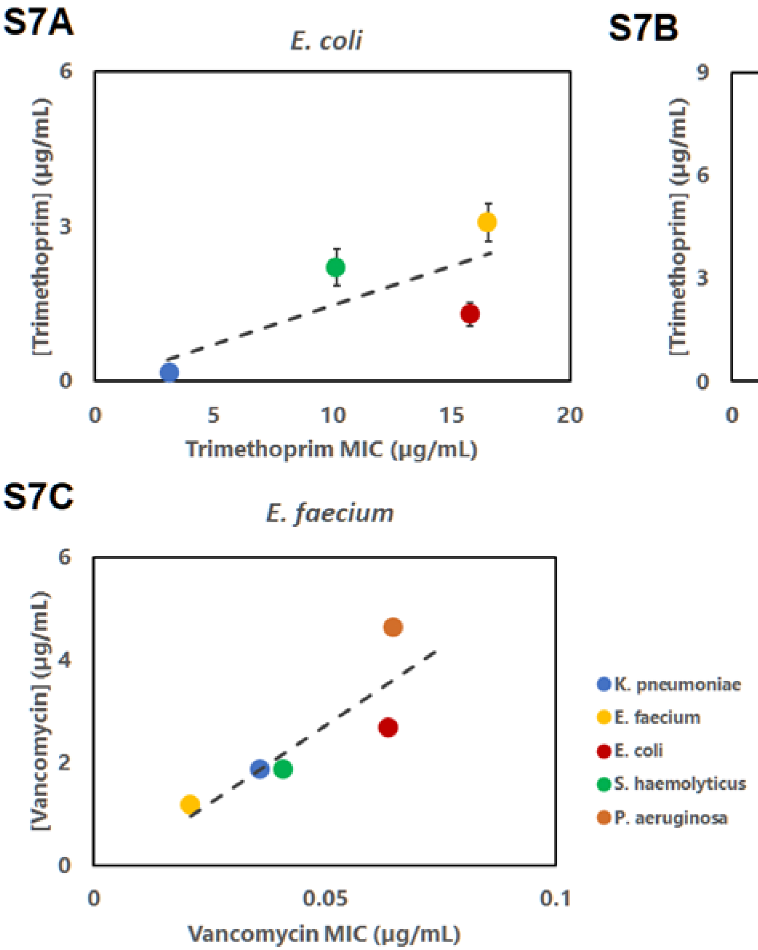

**Supplementary Figure 7 | Correlation of antibiotic concentration at the final transfer and MIC.** Antibiotic concentration at final transfer of the serial propagation experiment correlated with the MIC of evolved isolates in the absence of ecological interaction. Concentration sulfamethoxazole-trimethoprim is given per concentration trimethoprim, with ratio sulfamethoxazole:trimethoprim 5:1. Colors indicate the conditioned medium in which the mutants were evolved, with blue indicating *K. pneumoniae* conditioned medium, red *E. coli* conditioned medium, yellow *E. faecium* conditioned medium and green *S. haemolyticus* conditioned medium. The *E. faecium* mutants were also evolved in *P. aeruginosa* conditioned medium, represented by the orange marker. **A,** *E. coli* replicates show a low positive $R^2$=0.49, $P$<0.001, means ± s.e.m. **B,** *K. pneumoniae* replicates show a moderate negative $R^2$=0.68, $P$<0.001. **C,** *E. faecium* replicates show a high positive $R^2$=0.7297, $P$<0.001. For *K. pneumoniae* and *E. faecium* we find that the replicate lineages evolved in the same conditioned medium have identical MIC levels (B,C). For *E. coli* we observe a variability in the MICs, as we do for the replicate lineages in the evolution experiment (A, Figure 1A). Note that the end-point-antibiotic concentrations can deviate substantially from the MICs in the absence of these conditioned media. Particularly for *E. faecium*, the concentration of vancomycin at the final transfer is much higher (e.g., over 75 times, for *E. faecium* evolved in *P. aeruginosa* medium) than the MIC. This suggests that even though some antibiotic resistance has evolved, the tolerance or fitness altering effect conferred by the conditioned medium is important for enabling growth at these antibiotic concentrations.

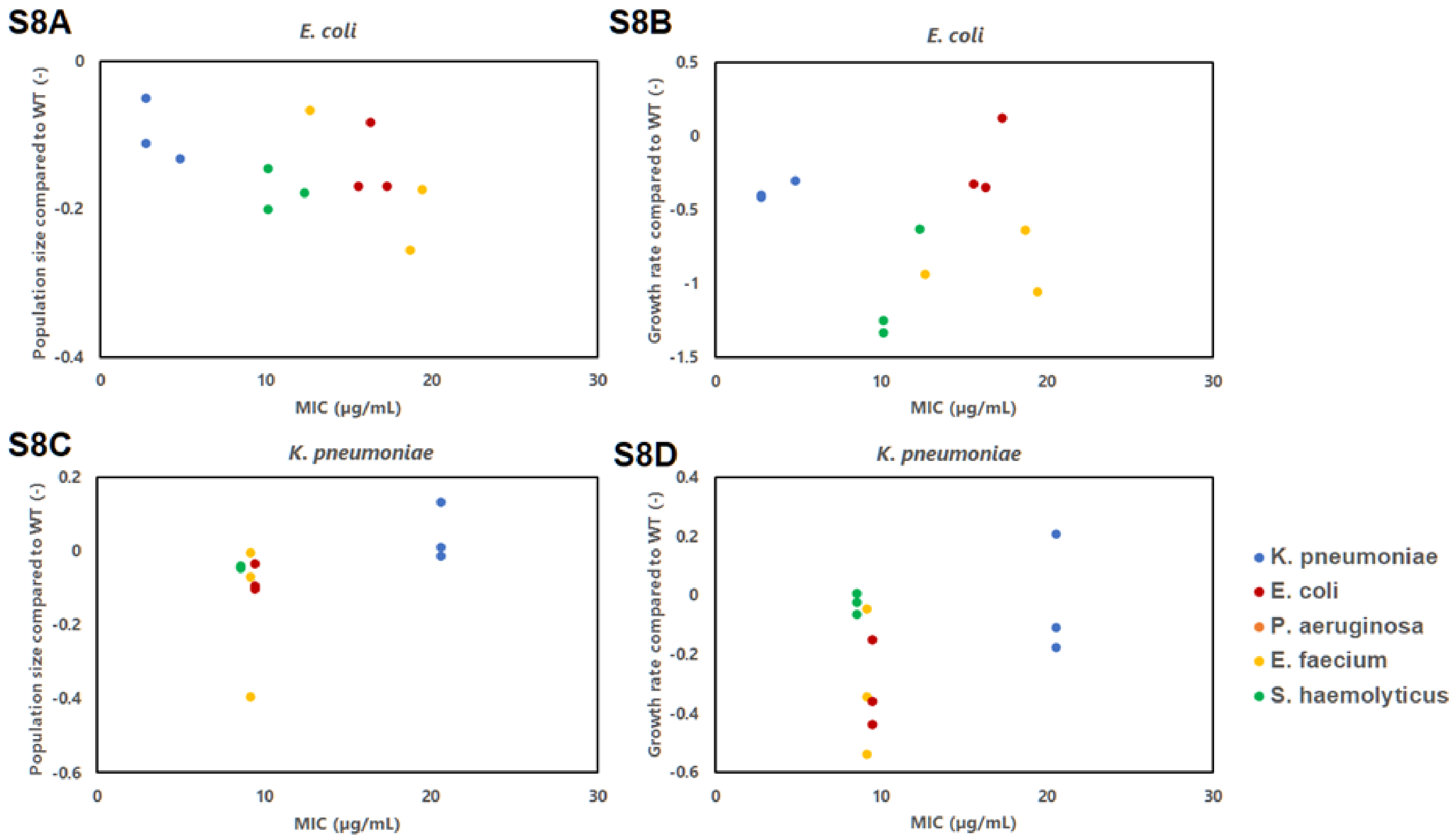

S8A
E. coli
Population size compared to WT (-)
MIC (µg/mL)
S8B
E. coli
Growth rate compared to WT (-)
MIC (µg/mL)
S8C
K. pneumoniae
Population size compared to WT (-)
MIC (µg/mL)
S8D
K. pneumoniae
Growth rate compared to WT (-)
MIC (µg/mL)
K. pneumoniae
E. coli
P. aeruginosa
E. faecium
S. haemolyticus

S8E S8F

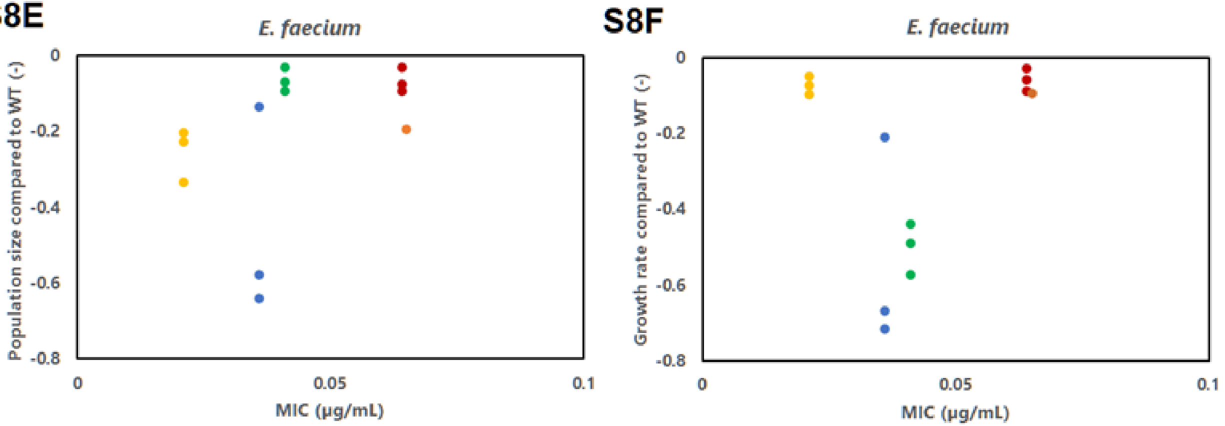

**Supplementary Figure 8 | Cost of resistance - Relationship between MIC and population size and growth rate of end-point clones in AUM.** Colors as shown in the legend represent the conditioned medium in which the replicates had evolved. MIC and growth were measured in 1x AUM, in the absence of microbial interactions. Replicate lineages cluster together for their MIC and cost of evolved resistance. **A** and **B** show the normalized population size and growth rate of *E. coli* replicates to the wild type *E. coli* respectively compared with the MIC to trimethoprim. $R^2$=0.27 and 0.0015, $P$=0.081 and 0.90 for the population size and growth rate respectively. **C** and **D** show the normalized population size and growth rate of *K. pneumoniae* replicates to the wild type *K. pneumoniae* respectively compared with the MIC to trimethoprim. $R^2$=0.24 and 0.14, $P$=0.11 and 0.23 for the population size and growth rate respectively. **E** and **F** show the normalized population size and growth rate of *E. faecium* replicates to the wild type *E. faecium* respectively compared with the MIC to vancomycin. $R^2$=0.17 and 0.045, $P$=0.16 and 0.49 for the population size and growth rate respectively.

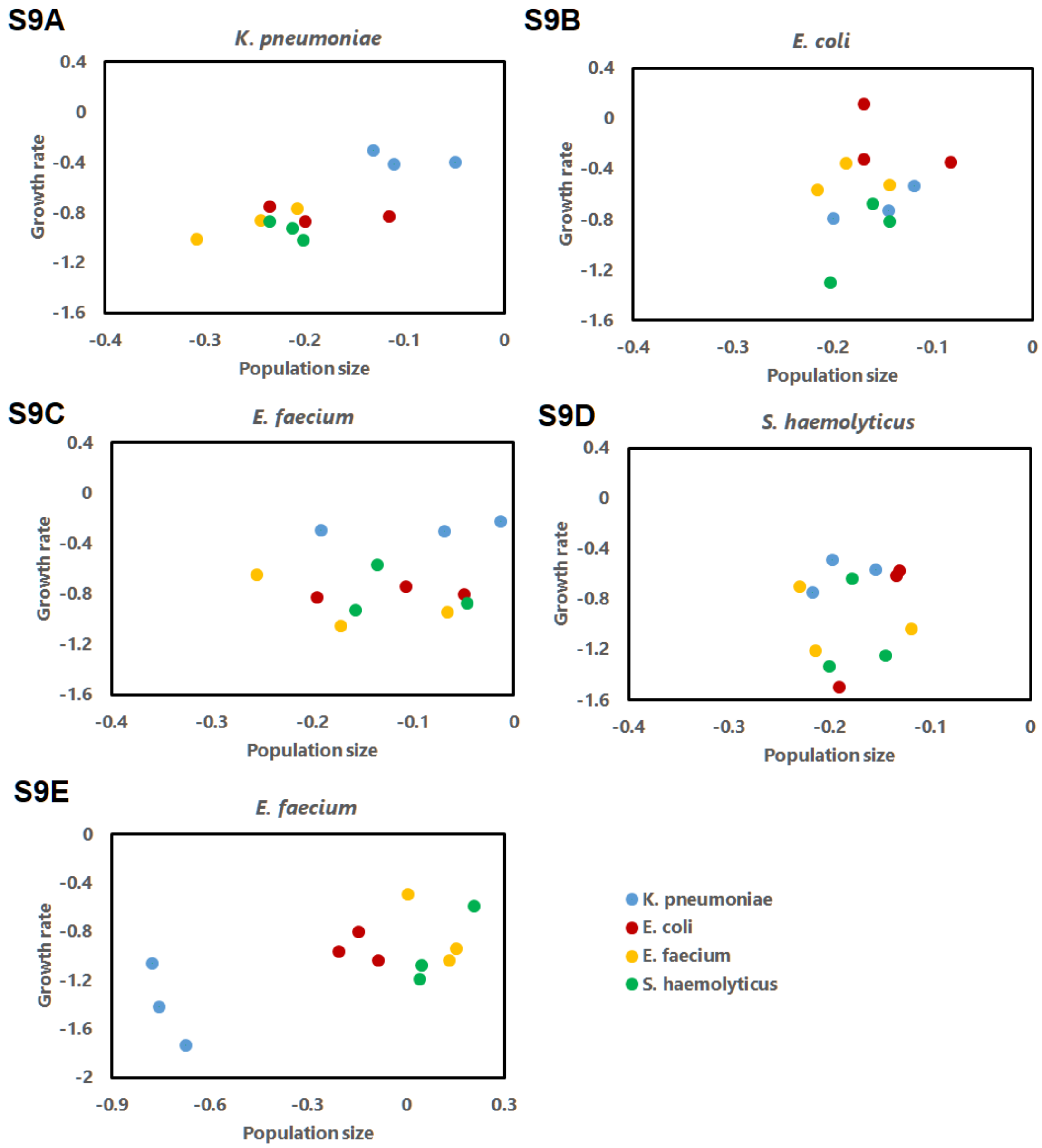

S9A
K. pneumoniae
Growth rate
Population size
S9B
E. coli
Growth rate
Population size
S9C
E. faecium
Growth rate
Population size
S9D
S. haemolyticus
Growth rate
Population size
S9E
E. faecium
Growth rate
Population size
K. pneumoniae
E. coli
E. faecium
S. haemolyticus

**Supplementary Figure 9 | Population size versus growth rate of evolved *E. coli* lineages.** Colors as shown in the legend represent the evolved conditioned medium background of the replicates. All data is normalized to the growth of the wild type in the respective conditioned medium, with a value of 0 indicating growth comparable to the wild type. Replicates were cultured in the different conditioned media environments as indicated above the figures, with **A** indicating *K. pneumoniae*, **B** *E. coli*, **C** *E. faecium*, and **D** *S. haemolyticus* conditioned media respectively. **E,** *E. coli* replicates cultured in *E. faecium* conditioned medium with antibiotics (0.5 µg/mL trimethoprim, 2.5 µg/mL sulfamethoxazole), normalized to wild type growth in regular AUM. Note that *E. coli* evolved in *E. faecium* conditioned medium grows quite poorly in that same *E. faecium* conditioned medium without antibiotics (Supplementary Figure 7C). *E. coli* lineages evolved in the other conditioned media even grow better in this environment. However, in the presence of antibiotics (Supplementary Figure 7E), these mutants outperformed the growth of the *E. coli* lineages in the other conditioned media. The cost and benefits of accrued mutations in these lineages seem to be dependent on both environmental factors, the presence of antibiotics ánd the conditioned medium.

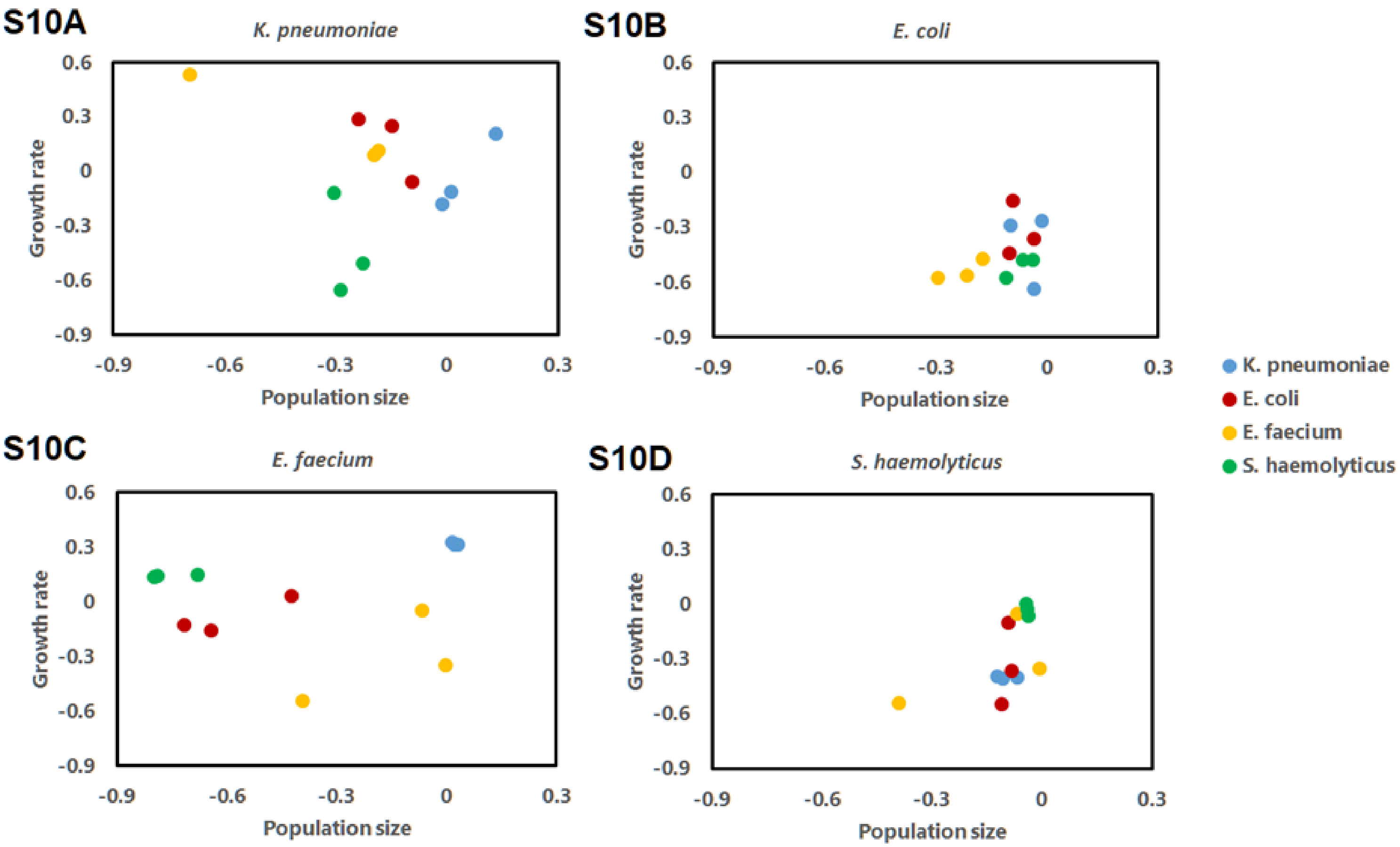

**Supplementary Figure 10 | Population size versus growth rate of evolved *K. pneumoniae* lineages.** Colors as shown in the legend represent the conditioned medium background of the replicates. All data is normalized to the growth of the wild type in the respective conditioned medium, with a value of 0 indicating growth comparable to the wild type. Replicates were cultured in the different conditioned media environments as indicated above the figures, with **A** indicating *K. pneumoniae*, **B** *E. coli*, **C** *E. faecium*, and **D** *S. haemolyticus* conditioned media respectively.

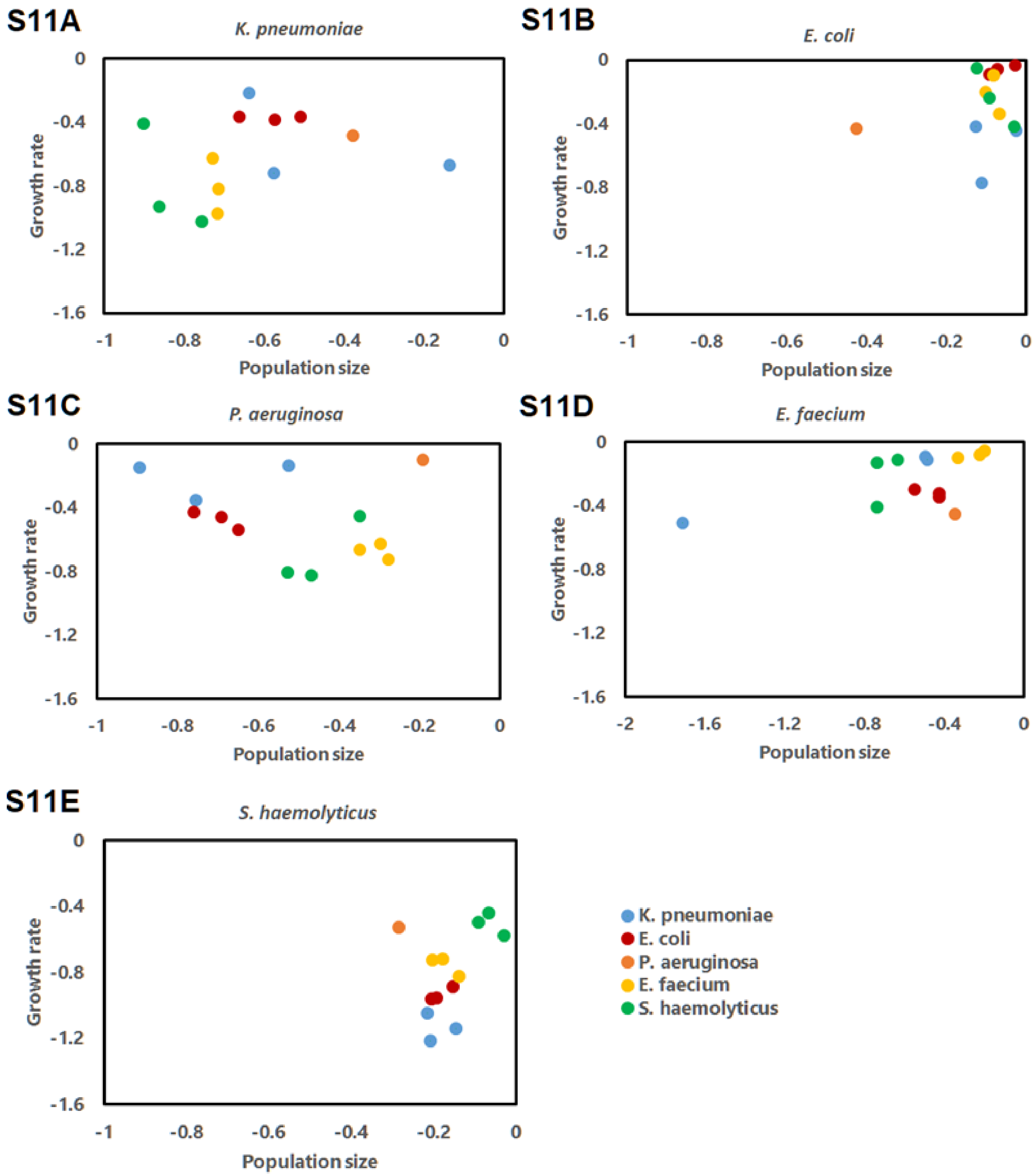

S11A
K. pneumoniae
Growth rate
Population size
0
-0.4
-0.8
-1.2
-1.6
-1
-0.8
-0.6
-0.4
-0.2
0
S11B
E. coli
Growth rate
Population size
S11C
P. aeruginosa
Growth rate
Population size
S11D
E. faecium
Growth rate
Population size
-2
-1.6
-1.2
-0.8
-0.4
S11E
S. haemolyticus
Growth rate
Population size
K. pneumoniae
E. coli
P. aeruginosa
E. faecium
S. haemolyticus

**Supplementary Figure 11 | Population size versus growth rate of evolved *E. faecium* lineages.** Colors as shown in the legend represent the conditioned medium background of the replicates. All data is normalized to the growth of the wild type in the respective conditioned medium, with a value of 0 indicating growth comparable to the wild type. Replicates were cultured in the different conditioned media environments as indicated above the figures, with **A** indicating *K. pneumoniae*, **B** *E. coli*, **C** *P. aeruginosa*, **D** *E. faecium* and **E** *S. haemolyticus* conditioned media respectively.

**Supplementary Table 1 | dN/ dS ratio focal isolates.**

| | ***E. coli*** | ***K. pneumoniae*** | ***E. faecium*** |
|---|---|---|---|
| Observed Nonsynonymous | 100 | 65 | 51 |
| Observed Synonymous | 22 | 9 | 3 |
| Observed dN / dS | 4.5455 | 7.2222 | 17 |
| Proportion Expected Nonsynonymous | 0.7286 | 0.7299 | 0.7518 |
| Expected dN / dS | 2.6848 | 2.7022 | 3.0294 |
| OB dNdS / EX dNdS | 1.6930 | 2.6727 | 5.6117 |
| Chisq | 5.12 | 8.27 | 10.74 |
| p-value | 0.024 | 0.004 | 0.001 |

**Supplementary Table 2 | Genes present on the different blocks of the two evolved plasmids in *K. pneumoniae* mutants.** Plasmid 1 corresponds to Supplementary Figure 1A and plasmid 2 corresponds to Supplementary Figure 1B.

| **Plasmid 1** | | | | | | | | | | | | | |
|---|---|---|---|---|---|---|---|---|---|---|---|---|---|
| **Block** | **Genes** | | | | | | | | | | | | |
| **1** | klcA | yubQ | traM | traY | traA | traL | traE | traK | traB | traP | traV | traC | |
| | trbI | traW | traU | trbC | traN | trbE | traF | traQ | trbB | traH | traG | traT | |
| | traD | traI | traX | finO | pld | repA2 | repA | repZ | repA4 | cdnC | cap6 | bcaP | |
| **2** | ydhP | cyp109 | TAH18 | cinB | yedK | resD | repE | parA | parB | samB | umuD | parM | yubD |
| | yubE | yubF | L7076 | yubI | ssb | yubM | arsH | arsC | arsB | arsR | ybaR | slr1230 | |
| **3** | crcB | tnpA | | | | | | | | | | | |
| **4** | gin | | | | | | | | | | | | |

| **Plasmid 2** | | | | | | | | | | | |
|---|---|---|---|---|---|---|---|---|---|---|---|
| **Block** | **Genes** | | | | | | | | | | |
| **1** | terA | uvrA | frmA | | | | | | | | |
| **2** | clpB | | | | | | | | | | |
| **3** | yjhV | insO2 | insN | PA0445 | intA | ImuB | JW5899 | vapC | Bmul_4720 | | |
| **4** | insB9 | insC | pcoS | pcoR | pcoD | pcoB | pcoA | silP | | | |
| | silA | silB | silC | fecR | fecA | fecB | fecC | fecE | | | |
| **5** | lacI | lacZ2 | lacY | pla | samB | sopB | sopA | repA | int | umuC | arsB |

**Supplementary Table 3 | List of mutated genes in *E. coli* mutants and corresponding proteins.** List corresponds to Figure 4A, genes that were found to be mutated in multiple final propagation mutants, with each unique mutation described.

| Gene | Protein | Mutation type | Length Indel (bp) | Mutation | Cluster of Orthologous Groups (COG) | COG category |
|---|---|---|---|---|---|---|
| *folA* | Dihydrofolate reductase | SNP | | Ile9Phe | Coenzyme Transport and metabolism | H |
| | | SNP | | N/A | | |
| | | SNP | | Thr25Met | | |
| | | INS | 1 | Tyr26fs | | |
| | | SNP | | Pro58Gln | | |
| | | SNP | | Asn60His | | |
| | | SNP | | Asn60His | | |
| | | SNP | | Asp64Glu | | |
| | | SNP | | Trp67Arg | | |
| | | SNP | | Ile131Leu | | |
| | | SNP | | Phe190Ser | | |
| *folM* | Dihydromonapterin reductase | DEL | 4 | Leu221fs | Coenzyme Transport and metabolism | H |
| | | DEL | 8395 | Asp65Gly | | |
| *folP* | Dihydropteroate synthase | SNP | | Pro64Leu | Coenzyme Transport and metabolism | H |
| | | SNP | | Pro64Ser | | |

| | | | | | | |
|---|---|---|---|---|---|---|
| | | SNP | | Thr62Ala | | |
| *nadR2* | DNA-binding transcriptional repressor/NMN adenylyltransferase | SNP<br>SNP<br>SNP<br>DEL | <br><br><br>23 | Pro151Leu<br>Pro151Leu<br>Ile211Asn<br>Ser259fs | Coenzyme Transport and metabolism | H |
| *acrR* | DNA-binding transcriptional repressor *acrR* | SNP<br>Upstream gene variant 1, SNP<br>Upstream gene variant 2, SNP<br>Upstream gene variant 3, SNP<br>Upstream gene variant 4, DEL | <br><br><br><br>1 | Thr32Ala<br>N/A<br>N/A<br>N/A<br>N/A | Transcription | K |
| *rpoC* | RNA polymerase subunit β' | SNP<br>SNP<br>SNP | | Ala328Pro<br>Glu1146Ala<br>Ile1357Ser | Transcription | K |
| *ycbZ* | Putative ATP-dependent protease *ycbZ* | Frameshift 1, DEL<br>Frameshift 2, DEL | 9<br>9 | N/A<br>N/A | Post-translational modification, protein turnover, chaperone functions | O |
| *deaD* | | SNP | | Arg624Gly | | L |

| | | | | | | |
|---|---|---|---|---|---|---|
| | ATP-dependent RNA helicase *deaD* | Frameshift 1, DEL | 124 | N/A | Replication, Recombination and Repair | |
| *dnaA* | Chromosomal replication initiator protein *dnaA* | SNP<br>SNP<br>SNP<br>SNP | | Glu361Gly<br>Ser228Phe<br>Arg87fs<br>Ile100Phe | Replication, Recombination and Repair | L |
| *phoQ* | Bifunctional sensor histidine kinase *phoQ* | SNP<br>SNP | | Gly384Ala<br>Ile207Asn | Signal Transduction Mechanisms | T |
| *rapZ* | RNase adaptor protein *rapZ* | SNP<br>SNP | | Leu144Gln<br>Ser22fs | Signal Transduction Mechanisms | T |
| *pabB* | Aminodeoxychorismate synthase subunit 1 | SNP<br>Upstream gene variant 1, SNP<br>Upstream gene variant 2, SNP | | Val8Met<br>N/A<br>N/A | Amino Acid metabolism and transport | E |
| *aroK* | Shikimate kinase 1 | SNP | | Pro121Leu | Amino Acid metabolism and transport | E |
| *adhE* | Fused acetaldehyde-CoA dehydrogenase and iron-dependent alcohol dehydrogenase dehydrogenase | SNP<br>SNP | | Ala626Val<br>Phe583Leu | Energy production and conversion | C |

| | | | | | | |
|---|---|---|---|---|---|---|
| *chbF* | Monoacetylchitobiose-6-phosphate hydrolase | INS<br>INS | 28<br>1 | N/A<br>N/A | Carbohydrate metabolism and transport | G |
| *gnd* | 6-phosphogluconate dehydrogenase, decarboxylating | SNP<br>Frameshift 1, INS | <br>1 | Gly143Arg<br>Ile6fs | Carbohydrate metabolism and transport | G |

**Supplementary Table 4 | List of mutated genes in *K. pneumoniae* mutants and corresponding proteins.** List corresponds to Figure 4B, genes that were found to be mutated in multiple final propagation mutants, with each unique mutation described.

| Gene | Protein | Mutation type | Length Indel (bp) | Mutation | Cluster of Orthologous Groups (COG) | COG category |
|---|---|---|---|---|---|---|
| *folA* | Dihydrofolate reductase | SNP<br>SNP<br>SNP<br>SNP<br>SNP<br>Upstream gene variant 1, SNP | | Ile94Leu<br>Trp30Arg<br>Leu28Arg<br>Asp27Glu<br>Met20Ile<br>N/A | Coenzyme Transport and metabolism | H |

| | | | | | | |
|---|---|---|---|---|---|---|
| | | Upstream gene variant 2, SNP | | N/A | | |
| *folM* | Dihydromonapterin reductase | SNP<br>DEL<br>SNP<br>SNP<br>SNP | <br>72 | Gly235Arg<br>Gene partially deleted<br>Ile150Phe<br>Ile102Asn<br>Ile11Ser | Coenzyme Transport and metabolism | H |
| *folX* | Dihydroneopterin triphosphate 2'-epimerase | SNP<br>SNP | | Ile100Ser<br>Ser111Leu | Coenzyme Transport and metabolism | H |
| *pCAR1* | IclR family transcriptional regulator, pca regulon regulatory protein | SNP<br>SNP<br>SNP<br>SNP<br>SNP<br>SNP<br>SNP | | Asp145Val<br>Ile139Ser<br>Arg138Gln<br>Thr122Pro<br>Glu116Ala<br>Asn115Ile<br>Leu114Phe | Transcription | K |
| *cspA* | Cold shock protein, *cspA* family | Upstream gene variant 1, SNP<br>Upstream gene variant 2, SNP | | N/A<br>N/A | Transcription | K |

| | | | | | | |
|---|---|---|---|---|---|---|
| | | Upstream gene variant 3, SNP | | N/A | | |
| | | Upstream gene variant 4, SNP | | N/A | | |
| *deoB* | Phosphopentomutase | SNP | | Asp326Ala | Carbohydrate metabolism and transport | G |
| | | DEL | 1 | Ala28fs | | |
| *pma1* | Plasma membrane ATPase | Upstream gene variant 1, SNP | | N/A | Inorganic ion transport and metabolism | P |
| | | Upstream gene variant 2, SNP | | N/A | | |
| *pitA* | Metal phosphate:H+ symporter *pitA* | SNP | | Gln364* | Inorganic ion transport and metabolism | P |
| | | INS | 50000 | N/A | | |

**Supplementary Table 5 | List of mutated genes in *E. faecium* mutants and corresponding proteins.** List corresponds to Figure 4C, genes that were found to be mutated in multiple final propagation mutants, with each unique mutation described.

| Gene | Protein | Mutation type | Length Indel (bp) | Mutation | Cluster of Orthologous Groups (COG) | COG category |
|---|---|---|---|---|---|---|
| *rpiR/CAC0191* | *rpiR*; Uncharacterized HTH-type transcriptional regulator *CAC0191* | SNP<br>SNP<br>SNP<br>SNP<br>INS | <br><br><br><br>1 | Glu68*<br>Ser156Ile<br>Gly223Ser<br>Arg244Gly<br>N/A | Transcription | K |
| *ompR* | Two-component system, o*mpR* family, response regulator | SNP | | Ter163* | Transcription/Signal Transduction Mechanisms | K T |
| *vanR* | Two-component system, *ompR* family, response regulator *vanR* | SNP<br>SNP<br>SNP<br>SNP<br>SNP<br>SNP<br>SNP<br>SNP | | Val13Ile<br>Glu23Lys<br>Gly24*<br>Asn104Lys<br>Ser114Phe<br>Asp177Tyr<br>Asp204Gly<br>Glu207Asp | Transcription/Signal Transduction Mechanisms | K T |

| | | | | | | |
|---|---|---|---|---|---|---|
| *spxA2* | Transcriptional regulator *Spx* | SNP<br>SNP<br>SNP<br>SNP | | Phe113Ser<br>Ile110Ser<br>Arg60Leu<br>Thr53Ala | Transcription/Signal Transduction Mechanisms | K T |
| *vanS* | Two-component system, *ompR* family, sensor histidine kinase *vanS* | SNP<br>SNP<br>SNP<br>SNP<br>SNP<br>SNP<br>SNP<br>SNP<br>SNP | | Ala28Glu<br>Phe44Leu<br>Asp65Glu<br>Ala119Ser<br>Asn169Asp<br>Gly186Glu<br>Thr203Met<br>Gly348Ser<br>Leu382Ile | Signal Transduction Mechanisms | T |
| *relA* | GDP/GTP pyrophosphokinase | SNP<br>DEL | <br>1 | Gln208*<br>Glu92fs | Signal Transduction Mechanisms/Nucleotide Transport and Metabolism | T F |
| *wecA* | Decaprenol-monophosphate-N-acetylglucosaminyltransferase | DEL<br>SNP | 10<br> | Ala460fs<br>Ala282Glu | Cell wall/membrane/envelope biogenesis | M |
| *yabM* | Putative membrane protein *yabM* | SNP<br>DEL | <br>4 | Gly12Val<br>Phe138fs | Cell wall/membrane/envelope biogenesis | M |

| | | | | | | |
|---|---|---|---|---|---|---|
| *nox* | NADH oxidase | Upstream gene variant 1, SNP<br>SNP<br>SNP | | N/A<br>Tyr31*<br>Arg430Cys | Secondary metabolites biosynthesis, transport and catabolism | Q |
| *prs2* | Phosphoribosylpyrophosphate synthetase | SNP<br>SNP<br>Upstream gene variant 1, SNP | | Lys186Asn<br>Pro147Thr<br>N/A | General Functional Prediction only | F |
| *pyk* | Pyruvate kinase | SNP<br>SNP | | Pro161Leu<br>Val259Phe | Carbohydrate metabolism and transport | G |
| *aroF* | Chorismate biosynthesis from 3-dehydroquinate | Complete DEL<br>DEL | 3459<br>1 | N/A<br>Lys202fs | Amino Acid metabolism and transport | E |
| *rex1* | Three prime repair exonuclease 1 | SNP<br>SNP<br>SNP | | Ala178Asp<br>Gly123Val<br>Ala97Val | Replication, Recombination and Repair | L |
| *hsdS* | Type I restriction enzyme EcoKI specificity protein | INV | | N/A | Defense machanisms | V |

**Supplementary Table 6 |** Intrinsic mutation rate of *K. pneumoniae* and *E. faecium*, measured as the phenotypic appearance of rifampicin resistant colonies in a mutation rate assay (Materials and Methods). The intrinsic mutation rates of *K. pneumoniae* and *E. faecium* are not significantly different (*t*-test, *N*=6, *P*=0.13).

| | **Average cfu /ml** | **Spots** | **Mutation rate** |
|---|---|---|---|
| ***K. pneumoniae*** | 6.44 | 10 | $1.86*10^{-08}$ |
| | | 8 | $1.47*10^{-08}$ |
| | | 11 | $2.05*10^{-08}$ |
| | | 10 | $1.86 *10^{-08}$ |
| | | 14 | $2.66 *10^{-08}$ |
| | | 16 | $3.08 *10^{-08}$ |
| **Average** | | | $1.86 *10^{-08}$ |
| | | **Spots** | **Mutation rate** |
| ***E. faecium*** | 2.28 | 5 | $2.54 *10^{-08}$ |
| | | 5 | $2.54 *10^{-08}$ |
| | | 7 | $3.60 *10^{-08}$ |
| | | 8 | $4.14 *10^{-08}$ |
| | | 12 | $6.35 *10^{-08}$ |
| | | 3 | $1.51 *10^{-08}$ |
| **Average** | | | $2.54 *10^{-08}$ |

**Supplementary information for *Microbial interactions affect the tempo and mode of antibiotic resistance evolution***

### Supplementary text 1

#### Additional information for the materials and methods

#### Bacterial isolates

Purity of pathogens was confirmed for the start of the evolution experiment by streaking on CHROMagar Orientation (CHROMagar) and selecting a single colony for overnight culturing in Lysogeny broth (LB) at 37°C. These cultures were turned into glycerol stocks (v/v 25% glycerol) and stored at -80°C.

#### Artificial urine medium

The 1x AUM medium was created by mixing 1.5 g/L bacto peptone L37 (BD), 3.15 g/L NaHCO3, 11.25 g/L urea (Roth), 4.8 g/L Na2SO4.10H2O, 1.8 g/L K2HPO4, 1.95 g/L NH4Cl, 15 mg/L bacto yeast extract (BD), 1.98 mM lactic acid (Roth), 600 mg/L citric acid, 105 mg/L uric acid, 1.2 g/L creatinine, 4.44 mg/L CaCl2.2H2O, 1.8 g/L Fe(II)SO4.7H2O (Riedel), 368 mg/L MgSO4.7H2O, 1.43 g/L KH2PO4. Chemicals were ordered from Sigma, unless stated otherwise. The pH was adjusted to 6.5 using 1M HCl.

**Preparation and replenishment of conditioned medium**

Bacterial isolates were cultured separately in 250mL AUM at 37°C and shaken at 200 rpm for 48h in 500mL Erlenmeyer flasks sealed using sterilized cotton wool and aluminum foil. Cultures were centrifuged at 4800xg for 15 minutes and pellets were discarded. Cell free supernatants were obtained by filtering the supernatants through a 0.45 µm and a 0.2 µm bottle top filter (TPP) in succession. The resulting cell free supernatant (or spent medium) was stored at room temperature in glass bottles. Conditioned medium was created by mixing 0.5 volume spent medium, with 0.17 volume 1x AUM and 0.33 volume 2.5x AUM (which only contained salt at 1x AUM concentration, to prevent excess salinity). The concentration in the final conditioned medium ranged from 0.75x AUM (for constituents that were consumed in the spent medium to 1.25x AUM (for constituents that were not consumed in the spent medium) (3). Any additional constituent or toxin produced by the donor species would be present at 0.5x medium concentration, due to the mixing with fresh media constituents.

**Serial dilution experimental evolution protocol**

The three focal isolates *K. pneumoniae*, *E. coli* and *E. faecium* were evolved in 5 ml conditioned medium, in 15 ml screw cap falcon tubes, with the caps slightly opened. The level of antibiotics at the start of the experiments was set at a 50% sub-MIC concentration (0.05:0.25 µg/mL trimethoprim:sulfamethoxazole for *K. pneumoniae,* 0.0125:0.0625 µg/mL trimethoprim:sulfamethoxazole for *E. coli* and 0.25 µg/mL vancomycin for *E. faecium*). In the absence of antibiotics, the stationary phase population sizes of *K. pneumoniae* and *E. coli* are ~$10^8$ CFU/ml, and *E. faecium* ~$10^4$ in 1x AUM. For each transfer, if OD600>0.1 for *K. pneumoniae* and *E. coli*, and OD600>0.05 for *E. faecium*, measured compared to reference, similar to McFarland standard, the antibiotic concentration was increased by 20%, otherwise the antibiotic concentration stayed at the same level as the previous transfer. Antibiotics were added to the conditioned medium just before each transfer of the focal species during the evolution experiment. Each transfer *K. pneumoniae* and *E. coli* accounted to maximally ~9 generations per bi-daily growth cycle ($2^9$=512), and *E. faecium* maximally ~6/7 generations per bi-daily growth cycle, $2^7$=128. Remaining cultures after transfer were stored, by preparing a 25% glycerol stock by mixing with 50% glycerol 1:1 v/v. Purity of the evolving cultures were checked on CHROMagar Orientation (CHROMagar) plates by plating 100µl directly from the cultures onto the plates and overnight incubation at 37°C. Because *K. pneumoniae* and *E. faecium* cultures had become contaminated after 60 days, the experiment was terminated earlier for those two pathogens.

### Antibiotic stocks

Antibiotic stocks were created separately by mixing 1 mg/mL trimethoprim in 96% ethanol, 5 mg/mL sulfamethoxazole in acetone and 5 mg/mL vancomycin in Milli-Q water. All antibiotics and solvents were ordered from Sigma. Stocks were stored at 4°C for up to two weeks and at -20°C for long term storage.

### 16S Phylogeny

A phylogenetic relationship between the isolates used in this study, was based on the 16S rDNA V1-V5 sequences (amplified with the 27F-16S, 926R-16S primers) obtained from de Vos, et al (3). The 16S sequences were aligned using Muscle multiple sequence alignment tool on the www.ebi.ac.uk website. The phylogenetic tree was constructed using the Clustal W tool on the same website.

### Growth measurements

Due to a -80°C freezer incident, only one lineage of *E. faecium* evolved in *P. aeruginosa* conditioned medium could be revived. Each isolate was incubated in three wells (triplicate cultures) of a preheated 96-well plate (transparent non-treated flat bottom, Brand) containing 200 μL of conditioned medium for 24 h. The plates were inoculated using a 96 solid pin multi-blot replicator tool, transferring ~1 μL of stock from a −80°C glycerol stock. The optical density at 600 nm (OD600) was measured every 6–7 min in the microplate reader (three flashes, 10 ms settle time). Before each measurement, the plate was shaken with a double orbital pattern. The population size of wild type cultures was obtained using the average OD600 at stationary growth after 24h. Evaporation of media from the plates was not detected. 96-well plates also contained non-inoculated wells as a control for contamination of both the conditioned medium and the inoculation step. Contamination was found to be negligible.

### MIC assays

Overnight cultures of *E. faecium, K. pneumoniae,* and *E. coli* grown at 37°C and agitated at 200rpm in AUM were transferred to fresh AUM, diluting the culture 100x. Cultures were allowed to grow for 6 h at 37°C, 200rpm to reach lag phase. Antibiotic discs were created by pipetting 10µL of desired concentration antibiotic on 6mm sterile Whatman antibiotic discs which were allowed to dry for 15 minutes. Antibiotic concentrations were calculated based on the concentration of the antibiotic in the final transfer multiplied by two.

MIC values were calculated from the inhibition zones observed on the agar plates after overnight growth. The time required for the antibiotic to diffuse through the agar and create the observed inhibition zone was estimated using its diffusion coefficient (4):

$$t = \frac{r^2}{2D} \qquad (1)$$

Where $t$ is the estimated time needed for the antibiotic to reach the radius $r$ of the inhibition zone in seconds, $r$ is the radius of the inhibition zone measured from the center of the antibiotic disc in cm, and $D$ is the diffusion coefficient of the antibiotic used in $cm^2/s$ which is assumed to be independent of the concentration. Diffusion coefficients for each antibiotic were calculated using the method described further down.

From there, the concentration of the antibiotic at radius $r$ ($C_{(r,t)}$) in µg/cm$^3$ can be calculated using (5):

$$C_{(r,t)} = \frac{A \cdot e^{(\frac{-r^2}{4Dt})}}{h4\pi Dt} \qquad (2)$$

Here, $A$ is the initial amount of antibiotics on the disc in µg, and $h$ is the height or thickness of the agar layer in cm.

Using the obtained concentration, the $MIC$ in µg/cm$^3$ of the bacterium can be calculated (6):

$$ln(MIC) = ln(C_{(r,t)}) - \frac{r^2}{4Dt} \qquad (3)$$

#### Method to obtain diffusion coefficients

Diffusion coefficients for each antibiotic were calculated assuming fluid dynamics in the agar. Viscosity and diffusion coefficients were obtained through the Stokes-Einstein equation (7):

$$D = \frac{k_b T}{6\pi\eta R} \qquad (4)$$

where $D$ is the diffusion coefficient in $cm^2/s$, $k_b$ is the Boltzmann's constant in $cm^2 \cdot g/s^2 \cdot K$, $T$ is the temperature in K, $\eta$ is the viscosity in g/cm·s, and $R$ is the radius of the molecule in cm. Following the known diffusion coefficient of fluorescein at 310.15 K in 1.5% agar of $8.5 \cdot 10^{-6}$, the viscosity of the agar can be estimated to be $4.31 \cdot 10^{-3}$ g/cm·s. The minimum size, or folded radius of the antibiotcs trimethoprim, sulfamethoxazole, and vancomycin were obtained from their respective molecular weight using the equation created by Erickson, 2009 (7):

$$R_{min} = 0.066 M^{\frac{1}{3}} \qquad (5)$$

With $R_{min}$ being the minimal radius of a sphere that can contain the mass of the folded molecule in nm and $M$ being the mass of the antibiotic in Da. Following this equation, we find the hydrodynamic radii of trimethoprim, sulfamethoxazole and vancomycin to be 0.44, 0.42 and 0.75 nm respectively.

#### Tolerance assays

We defined tolerance as the fold change difference of the level of an antibiotic that a population can sustain under the influence of bacterial interactions normalized to the level of antibiotic that a population can sustain in the absence of these interactions, as defined in (6). This is different from the definition of Brauner et al., that focuses on the time-dependence of tolerance. There, more tolerant strains of a bacterium have higher Minimum Duration for Killing (MDK), they describe that similar MICs do not necessarily mean similar tolerances (8).

### Sequencing

FASTQ read sequence files were generated using bcl2fastq2 version 2.18. Initial quality assessment was based on data passing the Illumina Chastity filtering. Subsequently, reads containing PhiX control signal were removed using an in-house filtering protocol. In addition, reads containing (partial) adapters were clipped (up to a minimum read length of 50 bp). The second quality assessment was based on the remaining reads using the FASTQC quality control tool version 0.11.5. The quality of Illumina reads was improved by trimming off low-quality bases using BBDuk, which is a part of the BBMap suite version 36.77. High-quality reads were assembled into contigs using ABySS version 2.0.2 (9). Scaffolding: The long reads were mapped to the draft assembly using BLASR version 1.3.1 (10). Based on these alignments, the contigs were linked together and placed into scaffolds. The orientation, order, and distance between the contigs were estimated using SSPACE-LongRead version 1.0 (10). Gap-closing and assembly polishing were performed using Illumina reads, gapped regions within scaffolds were (partially) closed using GapFiller version 1.10 (10). Finally, assembly errors and the nucleotide disagreements between the Illumina reads and scaffold sequences were corrected using Pilon version 1.21 (11).

### Detection of mutations and annotation

The raw reads were trimmed using TRIMMOMATIC (v 0.27, LEADING:3, TRAILING:3, SLIDINGWINDOW:4:15, MINLENGTH:70) (12). The trimmed reads were aligned on the reference genomes using bwa mem (v 0-7-15, default parameters). After mapping reads were filtered for quality and sorted using samtools (v 0.1.19) (13). Duplicates were removed using picard tools (v.2.8.2) and realignment around indels was performed with GATK (v .3.7-0) (14).The resulting BAM files were combined in mpileup format (samtools, mpileup, default parameters, v 0.1.19) (13), after which SNPs and indels were separately called using varscan (v 2.3.9, using mpileup2snps and mpileup2indel resp., --output-vcf 1, --min-var-freq 0.05) (15) for which we only kept those variants that differed between the samples. The resulting raw vcf files was then annotated with snpEff (v 4.3) (16) using the annotation file (GFF) as input. For structural variant detection we followed a split- and clipped-read based method (17).

The resulting vcf files were further filtered using vcfR (v 1.8.0) (18). Using vcfR the number of alternative reads (non-reference) were divided by the total read depth to obtain the alternative read frequency. Variants with a frequency between 10 and 90% in any individual were omitted. These values leave some margin for sequencing and mapping errors but leave out variants with too many. For a variant an individual had to have at least 10x coverage to be called. Variants were identified as different from the ancestor if this frequency differed by more than 80%. In most cases sampled different by 100% from the ancestor, but in cases where both ancestor and evolved strains showed sequencing error, this frequency can deviate from 100%. Initially all variants were omitted if they had more than 30 NA calls, meaning that 30 samples lacked any coverage. However, all variants showed enough coverage to be called in the final variant set. All variants identified were visually inspected using IGV (19) (v 2.8.0).

## dN/dS

We calculated the observed dN/dS by dividing the number of nonsynonymous substitutions with the synonymous substitutions in the genomes of each focal species. We calculated the expected proportion of nonsynonymous versus synonymous in the following way. We theoretically mutated all codons, controlling for mutation bias extracted from Hershberg & Petrov (20) to extract the proportion of nonsynonymous versus synonymous substitutions. For ATG for instance this is 1:0 because all changes lead to a nonsynonymous change, while for GCA this is for instance 2/3. Then we used the predicted whole genome coding sequence from the newly assembled reference genomes of each ancestor to calculate the whole genome codon usage. Multiplying the proportionate codon usage with the probability of finding a nonsynonymous substitution then gave us the expected proportion of a nonsynonymous substitution per genome (see Supplementary table 6). This gave us the expected values of dN/dS which were 2.7 for *K. pneumoniae* and *E. coli*, while it was 3.0 for *E. faecium*. Using a chi square test, we calculated a significant deviation from this value using the observed 7.2, 17 and 4.5 dN/dS for *K. pneumoniae*, *E. faecium* and *E. coli* respectively.

### Mutation rate assay

To estimate the mutation rate, we used a high-throughput test which is similar to the fluctuation test developed by Luria and Delbrück to test the mutation rates in bacterial population(21). A pre-culture of each focal species was made in LB, and diluted in pre-warmed LB to a density of OD600 $10^{-3}$. 23 µl of this diluted culture was added to 23 ml of LB or conditioned medium (inoculated density OD600 $10^{-6}$). Six 96-well plates were filled with 200 µl of this cell-medium mixture in each well. The 96-well plates were incubated overnight at, at static, non-shaking conditions. After 16 hours, the plates were placed in BMG Clariostar plate reader, and mixed for 240 seconds at 600rpm, double orbital, and the $OD_{600}$ was recorded. Following overnight growth, 8 µl drops from each well were spotted on pre-warmed LB agar plates containing rifampicin (*E. faecium* 4 µg/ml MIC, 15 µg/ml used in test, *K. pneumoniae* 40 µg/ml MIC, 120 µg/ml used in test). After two days of incubation at 37°C, the number of spots that contained at least one antibiotic-resistant colony, were counted. We did not obtain any mutation rates of *E. coli*, because this isolate was highly resistant to rifampicin (>500 µg/ml).

The mutation rate was estimated using, $p_0=e^{-m}$, where $m=a(N_t-N_0)$ (21). $p_0$ is the change of a culture not resulting in a spot with a resistant mutant. *m* is the number of mutations that have occurred during a specific time interval. In this case this is the overnight growth where the very small starting population, *N0*, has given rise to the final population, *Nt*, over the course of several generations during which the mutations could have occurred. This translates into the mutation rate, *a*. In other words, the number of new cells ($N_t$-$N_0$) times the mutation rate (*a*) gives the number of mutations (*m*) for a specific site on the genome which is inversely related to the chance of NOT getting a resistant cell in the culture (*p0*). a is estimated using the following formula: *a*= log(1/(1-(spot-count/96)))/[*CFU/ml],* where 1-(spot-count/96)= $p_0$.

#### *H*-index databases

Mutated genes were categorized in their corresponding COGs using the COG database of the National Centre for Biotechnology Information (NCBI) (22), with cross-referencing to Uniprot database (23).

A thorough schematic explanation of the H-index calculation is given in Supplementary Figure 9 of Schenk, *et al* (24).

#### Statistical analyses

Throughout all of the analyses, whenever the values between two conditions of mutants were compared, a two-tailed Welch's t-test was performed to test the null hypothesis that the two parameters do not have significantly different values (hypothesis of neutrality). We used a Poisson-test to assess whether the number of mutations in the mutators was significantly different in the hypermutators and the non-hypermutators. We used the square of Pearson correlation coefficients and corresponding *P*-values to determine the level of correlation. We used a Kruskal-Wallis rank sum test, followed by a Chi-Squared test, to determine whether particular conditioned media were significantly overrepresented for their effect of the speed up or slowdown of evolution. To compare whether the combination of population size and growth rates of mutants from a specific conditioned medium background were different from those of other backgrounds, we used a one-way ANOVA with post-hoc Tukey’s test for all combinations.

### Supplementary text 2

#### Specific mutations within evolved lineages

If we zoom in on the specific mutations in our replicate lineages for the dihydrofolate reductase gene *folA*, we find that most of the non-synonymous mutations in the *folA* gene in *K. pneumoniae* and *E. coli* have been described before. The L28R mutation in the coding region of *folA*, was detected in *K. pneumoniae* evolved in the conditioned medium of *E. coli* and *S. haemolyticus*. This specific mutation was hypothesized to be favored under strong selective conditions previously (25). *K. pneumoniae* evolved in *K. pneumoniae* and *E. faecium* conditioned media had a more diverse spectrum of *folA* mutants, consistent with weaker selective conditions (25). Indeed, these lineages have a slower resistance trajectory compared to *E. coli* and *S. haemolyticus* (Figure 1B). Mutations D27E, W30R, and I94L, are known to decrease trimethoprim binding and are generally found in *E. coli* mutants that were evolved in all conditioned media, yet specific to *K. pneumoniae* mutants evolved in *E. faecium* and *K. pneumoniae* conditioned media (25). *E. coli* mutations I5F, P21Q, and F153S, also lead to decreased binding of the antibiotic. The P21Q mutation was found in mutants evolved in all conditioned media, and has been described as a common mutation in both clinical and laboratory setting (25,26). Yet the mutation N23H in *E. coli* mutants evolved in *E. coli* and *S. haemolyticus* conditioned media, has not been described before in *E. coli* (27). Yet, the N23 residue is located near the binding site of folate and cofactor (28).

The specific mutations of the dihydropteroate synthase gene *folP* have been observed before in other studies (29). The P64S mutation lies in a highly conserved region that is linked to a strong increase in resistance to sulfonamide antimicrobial agents, such as sulfamethoxazole. The proximity of P64 to R63 of dihydropteroate synthase where sulfamethoxazole binds through a hydrogen bond likely altered the binding of sulfamethoxazole to the enzyme (29).

Stress response genes are also implicated to be involved in antibiotic resistance (30). The *cspA* gene in *K. pneumoniae* and the *deaD* gene in *E. coli* are both involved in a general stress response (cold-shock) (31–33). The *cspA* gene was targeted in *K. pneumoniae* in the media of *E. coli* and *S. haemolyticus* and *deaD* was targeted in *E. coli* in *E. coli* conditioned medium respectively (34) (Figure 5A,B).

Additionally, efflux seems to play a role in the resistance evolution of *E. coli*. Two genes involved in the AcrAB-TolC multidrug efflux pump (35), *acrR* and *phoQ*, were targeted in *E. coli* mutants evolved in the Gram-positive *E. faecium* and *S. haemolyticus* conditioned environments. Other mutations related to efflux were detected in one *E. coli* lineage that evolved in *E. faecium* conditioned medium. Duplication of a small part of the *marR* (36) promoter region potentially led to altered cis-regulatory effects of the marRAB operon. A lower expression of *marR* leads to the upregulation of efflux and the downregulating of influx mediated by *ompF* (37,38). These efflux related mutations occurred in the fast-evolving lineages with the largest initial population sizes (Figure 2A), and therefore do not correlate with smaller population sizes and potentially weaker selection as observed in other studies (39). This suggests that the fitness effects of antibiotic resistance mutations for *E. coli* and *K. pneumoniae* are conditioned medium specific, instead of governed by the supply of the same set of mutations via effects on population size.

Also, different genes involved in redox homeostasis and the regulation of oxidative stress are targeted in *E. faecium* in the presence of the different bacterial exudates. Specifically, in *K. pneumoniae* and *E. faecium* conditioned medium *nox1* was targeted, whereas in *E. coli*, *P. aeruginosa* and *S. haemolyticus* medium the transcriptional regulator *rex1* was hit. The *Nox1* gene encodes a NADH oxidase (40,41), and *rex1* is involved in the control of NADH/NAD(+) levels (42). The *spxA* gene was target in the conditioned medium of *P. aeruginosa* and *E. faecium*, and is also related to the oxidative stress tolerance (43). In addition, in *P. aeruginosa* conditioned medium *relA*, involved in the stringent response, was targeted. This target was previously found to be associated with multi antibiotic tolerance *in vivo*(44).

Concluding, specific clinically relevant antibiotic resistance related genes were targeted more than once in parallelly evolved populations in conditioned medium (Figure 5), this shows that genetic basis of antibiotic resistance evolution is mediated by the genetic background of the focal species, as well as on the ecological interactions.
